# Immune receptor LILRB1 mediates *cis*-signalling which is targeted by RIFINs of the malaria parasite

**DOI:** 10.64898/2026.04.16.718981

**Authors:** Samuel G. Chamberlain, Marcus A. Widdess, Alexander M. Mørch, Akihito Sakoguchi, Rina Sakuno, Elke Kurz, Lina Chen, Salvatore Valvo, Shiroh Iwanaga, Michael L. Dustin, Matthew K. Higgins

## Abstract

The malaria parasite, *Plasmodium falciparum*, replicates within human erythrocytes, where it is susceptible to clearance by immune cells. It uses erythrocyte surface proteins, the RIFINs, to signal through immune receptors and to suppress immune cell function. Previous studies identified two groups of RIFINs which bind different sites on inhibitory immune receptor LILRB1. While some RIFINs bind an elongated LILRB1 conformation, triggering inhibitory immune signalling in ‘*trans’*, we show that other RIFINs stabilise a c-shaped LILRB1 conformation. This buckled LILRB1 binds MHC class I found on the same immune cell, which triggers inhibitory ‘*cis’* signalling. Therefore, LILRB1 exists in dynamic equilibrium, with an elongated conformation able to bind to ligands in ‘*trans*’ on a target cell, while a buckled conformation signals through MHC class I in ‘*cis*’, setting the signalling threshold. Different clades of RIFINs exist to mediate inhibitory signalling through each of these LILRB1 conformations to prevent parasite destruction.

## Introduction

The deadliest form of malaria is caused by the parasite, *Plasmodium falciparum*, with the highest mortality in African children under 5 years old. The clinical symptoms of malaria occur during the blood stage of infection when parasites replicate within red blood cells. To prevent splenic clearance, the parasites place PfEMP1 adhesion proteins onto infected RBC (iRBC) surfaces. These PfEMP1 bind human endothelial receptors, leading to sequestration of iRBCs to blood vessel and tissue surfaces, away from the spleen ^1,2^. The PfEMP1 have diversified into a large protein family, allowing antigenic variation, but remain a target for antibody-mediated detection. Antibodies that target PfEMP1 are found in patient sera, increase with age, and correlate with decreasing susceptibility to life-threatening malaria symptoms ^3,4^. Indeed, antibody-dependent cellular cytotoxicity (ADCC) by natural killer (NK) cells or γδ T lymphocytes contribute to parasite suppression and clearance ^5–7^.

Members of a second protein family, the RIFINs, are also displayed on iRBC surfaces^2^. Multiple different RIFINs have recently been shown to bind different human inhibitory immune receptors, including LILRB1 ^8^, LILRB2 ^9^, LAIR1 ^10^ and KIRs ^11^. This localises these inhibitory receptors to immune synapses, triggering inhibitory signalling and suppressing immune cell function. Several adaptions of the human immune system have most likely occurred in response to RIFINs. First, remarkable antibodies have evolved which contain the ectodomains of LAIR1 and LILRB1 inserted into their Fab regions^12^. Second, RIFINs have been identified that bind the activating KIR, KIR2DS1, activating NK cells and increasing killing ^11^. This raises the intriguing possibility that activating KIRs have evolved, in part, to counter triggering of inhibitory pathways by RIFINs that bind inhibitory KIRs ^13,14^.

LILRB1 was the first immune receptor shown to be regulated by RIFINs ^8^. It is an inhibitory receptor expressed on the surfaces of γδ T lymphocytes, NK cells, and monocytes^15,16^. LILRB1 binds MHC class I in trans, triggering a ‘don’t kill me’ signal that reduces immune cell activation ^16^. LILRB1 consists of an ectodomain with four immunoglobulin domains (D1-D4) arranged as two didomains (D1D2 and D3D4) and an intracellular ITIM sequence, associated with inhibitory signalling. Multiple LILRB1-binding RIFINs bind the D2 domain of LILRB1, overlapping with the MHC class I binding site ^17^. This molecular mimicry allows the RIFINs to elicit the same inhibitory signal as MHC class I, supressing perforin release from NK cells ^17^. More recently, antibodies were discovered in malaria-exposed Malian volunteers which contained inserts of the D3 and D4 domains of LILRB1 in the elbow of the heavy chain of the Fab region. These antibodies, such as MDB1, were also found to bind to multiple RIFINs, which are different from those which bind LILRB1 D2 domain ^12^. This suggested that there are two groups of LILRB1-binding RIFINs, one which binds to the D1D2 didomain and a second which binds the D3D4 didomain. We therefore aimed to determine whether RIFINs which bind to D3D4 domains of LILRB1 also mediate inhibitory signalling in NK cells and to understand why the parasite has evolved two distinct groups of RIFINs which target different regions of the same immune receptor.

## Results

### A structure of LILRB1 in complex with a RIFIN reveals a c-shaped LILRB1 conformation

Atypical antibodies have been identified in adults in a malaria-endemic region in which domains 3 and 4 (D3D4) of the inhibitory immune receptor LILRB1 have been directly inserted into their heavy chain Fab fragment ^12^. The structure of a Fab fragment of one of these antibodies (MDB1) bound to a *Plasmodium falciparum* RIFIN, *Pf*3D7_1373400 (D3-RIF), revealed the RIFIN to bind a hydrophobic pocket on the D3 domain of LILRB1 (Figure 1a). Superimposition of this structure onto the known structure of LILRB1, which adopts an extended conformation^17,18^, revealed extensive clashes between the RIFIN and the site of inter-domain contact between the D2 and D3 domains of LILRB1 (Figure 1a). This raised the question of whether this interaction, which has been observed in antibodies lacking the first two domains of LILRB1, can also occur with full-length LILRB1 and what consequences this might have for LILRB1 conformation.

**Figure 1:**
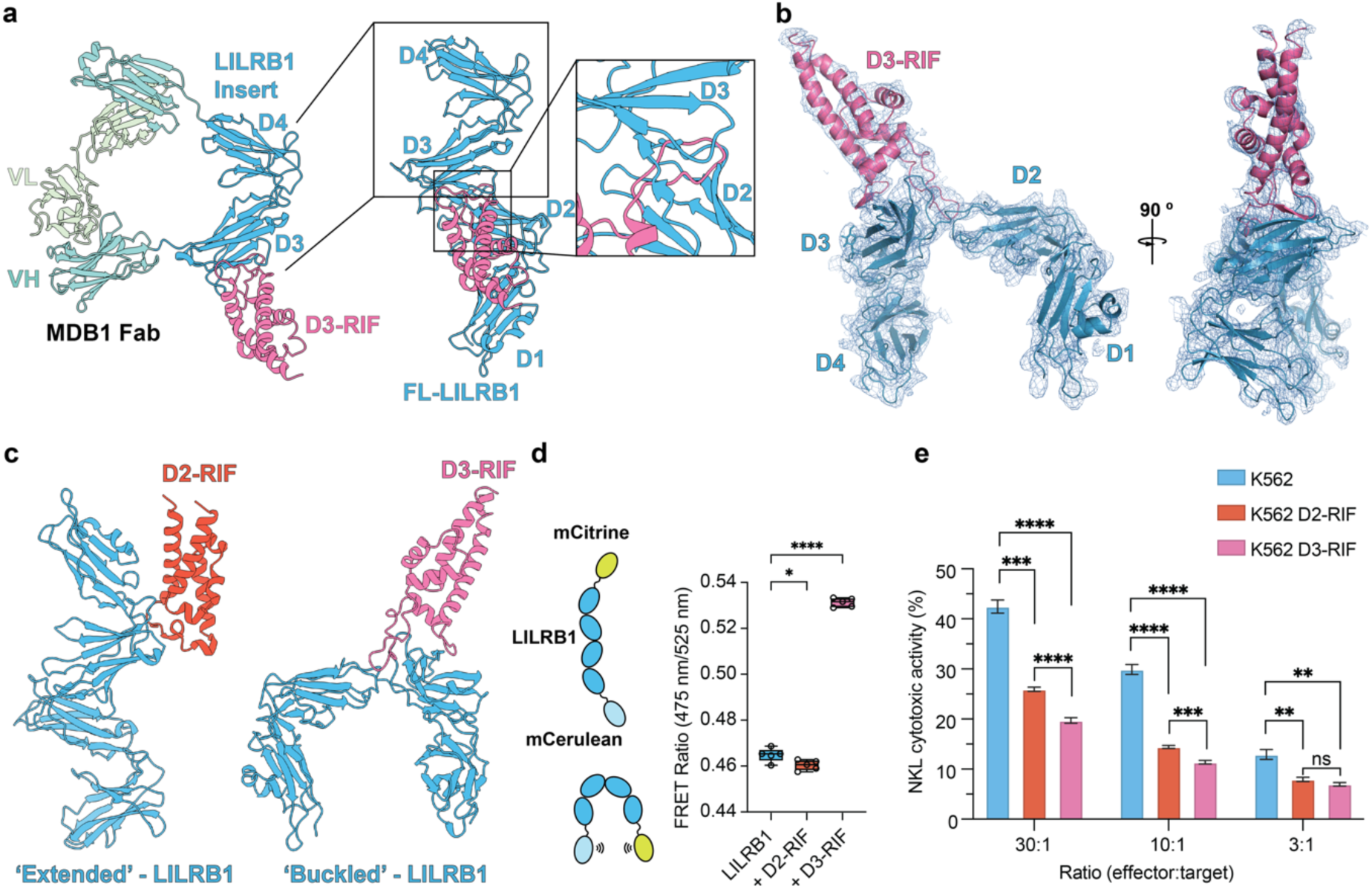
LILRB1 adopts a buckled conformation to bind D3-RIF, *Pf*3D7_1373400. **a)** The structure of antibody MDB1 (PDB code 7KFK) bound to RIFIN D3-RIF (left). Domains 3 and 4 (D3D4) of LILRB1 (blue) are inserted in the hinge between the VH region of the Fab (VH in teal, VL in light green) and D3-RIF (pink) binds the D3 domain. This RIFIN binding site is incompatible with all current structures of full-length (FL) LILRB1, as shown by superimposing D3-bound D3-RIF on the structure of elongated LILRB1 (centre, PDB code 6AEE), due to a large clash between the ⍺3-4 binding loop of D3-RIF and where the D2 domain of LILRB1 contacts the D3 domain (right). **b)** The structure of D3-RIF (pink) in complex with LILRB1 (blue), shown in cartoon representation with blue mesh denoting the experimental electron density. **c)** Comparison of the structures of D2- (orange) vs D3-RIF (pink) bound to LILRB1 (blue). **d)** Schematic representation of the LILRB1-FRET sensor reporting on the conformation of LILRB1 (left) and FRET ratio (475 nm/525 nm) of the LILRB1 FRET probe after excitation at 433 nm either alone (1 µM) or in the presence of D2-and D3-RIFs (10-fold excess) (right). Datapoints represent measurements from five independent experiments and two biologically independent preparations of the protein. Statistical significance was determined by one-way ANOVA in PRISM10. *p <0.05, **** <0.0001. **e)** Effect of K562 cells expressing D3-Rif on the cytotoxicity of NKL cells tested at three ratios of target to effector cells. Data represent the mean± s.d. (n=4 biologically independent samples) with ****P < 0.0001, ***P < 0.001, **P < 0.01 (two-sided Student’s t-test).

We therefore crystallised a complex of D3-RIF (residues 173 – 334) and the LILRB1 ectodomain. Repeated iterations of screening and optimization yielded crystals that diffracted to 4.8 Å, with the resolution likely limited by LILRB1 flexibility. Molecular replacement using structures of D3-RIF and two LILRB1 di-domains (D1D2 and D3D4) allowed structure determination, with just one copy of the complex in the asymmetric unit (Figure 1b, Table S1). Structures of individual components were docked into the electron density and refined to reveal the architecture of the complex. The interface between the RIFIN and the D3D4 domain of LILRB1 was very similar to that previous observed ^12^ (Supplementary Figure 1a) ^19^. However, in contrast to previous extended structures of LILRB1, alone or bound to a RIFIN which contacts the D2 domain of LILRB1, LILRB1 bound to D3-RIF adopted a c-shaped conformation due to extensive bending at the hinge between domains 2 and 3, bringing domains 1 and 4 close together (Figure 1c). This allows the D3 RIFIN to occupy the hydrophobic pocket normally involved in D2-D3 contacts.

To determine whether the buckled conformation of LILRB1 also occurs outside the crystal environment a FRET probe was designed, in which the LILRB1 ectodomain was flanked with mCerulean and mCitrine ^[19]^. After confirming that there were no concentration dependent auto-FRET effects, spectral scans were carried out by exciting at 433 nm and measuring the emission ratio between 475 nm and 525 nm (Figure 1d, Supplementary Figure 1b,c). Measurements were then recorded for the FRET sensor alone or in the presence of either the D2-binding RIFIN, *Pf*3D7_1254800 (D2-RIF) or D3-RIF (Figure 1d). The FRET ratio increased significantly in the presence of D3-RIF but not D2-RIF. This is consistent with the N- and the C-terminal fluorescent tags generally being apart, suggesting that LILRB1 favours an extended conformation, but which are then brought into closer proximity due to buckling of the LILRB1 structure, which is stabilised by D3-RIF. Indeed, measurements at different concentrations of D3-RIF showed that a dose dependent effect can be observed which fitted to a 1:1 binding model (PRISM one-site total) with an EC_50_ close to the K_D_ of D3-RIF for LILRB1 (Supplementary Figure 1d). These data show that LILRB1 in solution is found predominantly in an extended conformation, with a buckled confirmation stabilised by binding of the D3-RIF.

### D2- and D3-LILRB1 binding RIFINs supress NK function

Previous work has shown that multiple RIFINs that bind D1D2 of LILRB1 can suppress NK cell activation ^8,17^. We therefore asked whether a RIFIN that binds the D3 domain of LILRB1 also triggers inhibitory signalling. To test this, the NK cell line, NKL was incubated with target K562 cells either expressing D2-RIF or D3-RIF. At three different effector-target ratios, both D2- and D3-binding RIFINs exhibit a suppressive effect on NKL killing (Figure 1e). Indeed, D3-RIF suppressed NKL cell function to a greater degree than D2-RIF. Therefore, a RIFIN which binds only to the buckled conformation of LILRB1 can increase inhibitory signalling.

### LILRB1-D3 binding RIFINs are common in multiple P. falciparum genomes and all bind the buckled LILRB1 conformation

A previous study identified seven RIFINs in the 3D7 genome that bind LILRB1 D3D4-containing antibodies ^12^. We asked whether this represents the full repertoire of LILRB1 D3D4-binding RIFINs from 3D7 by screening a library of infected red blood cells (iRBCs), each expressing a single RIFIN ^8,11^. Two biologically independent RIFIN libraries were transfected into iRBCs (rif-lib1 with 98.0% RIFIN coverage (154/157), rif_lib2 with 97.5% coverage (153/157)), both of which showed high RIFIN coverage by amplicon-seq. Flow sorting with LILRB1 D3D4-Fc identified 16.3% and 15.6% of positive cells in rif-lib1 and rif-lib2, respectively and these were returned to culture for enrichment (Figure 2a and Supplementary Figure 2a). Enriched iRBC libraries (47.4% and 47.7%, rif-lib1 and rif-lib2) were subjected to amplicon-seq, identifying 20 and 21 RIFINs with 18 of these shared between the two biologically independent repeats. Six out of the seven previously identified RIFINs ^12^ were present in our candidate list and 11 of these candidates clustered into one clade (Figure 2b, Supplementary Figures 2b and 3). ^[12]^

**Figure 2:**
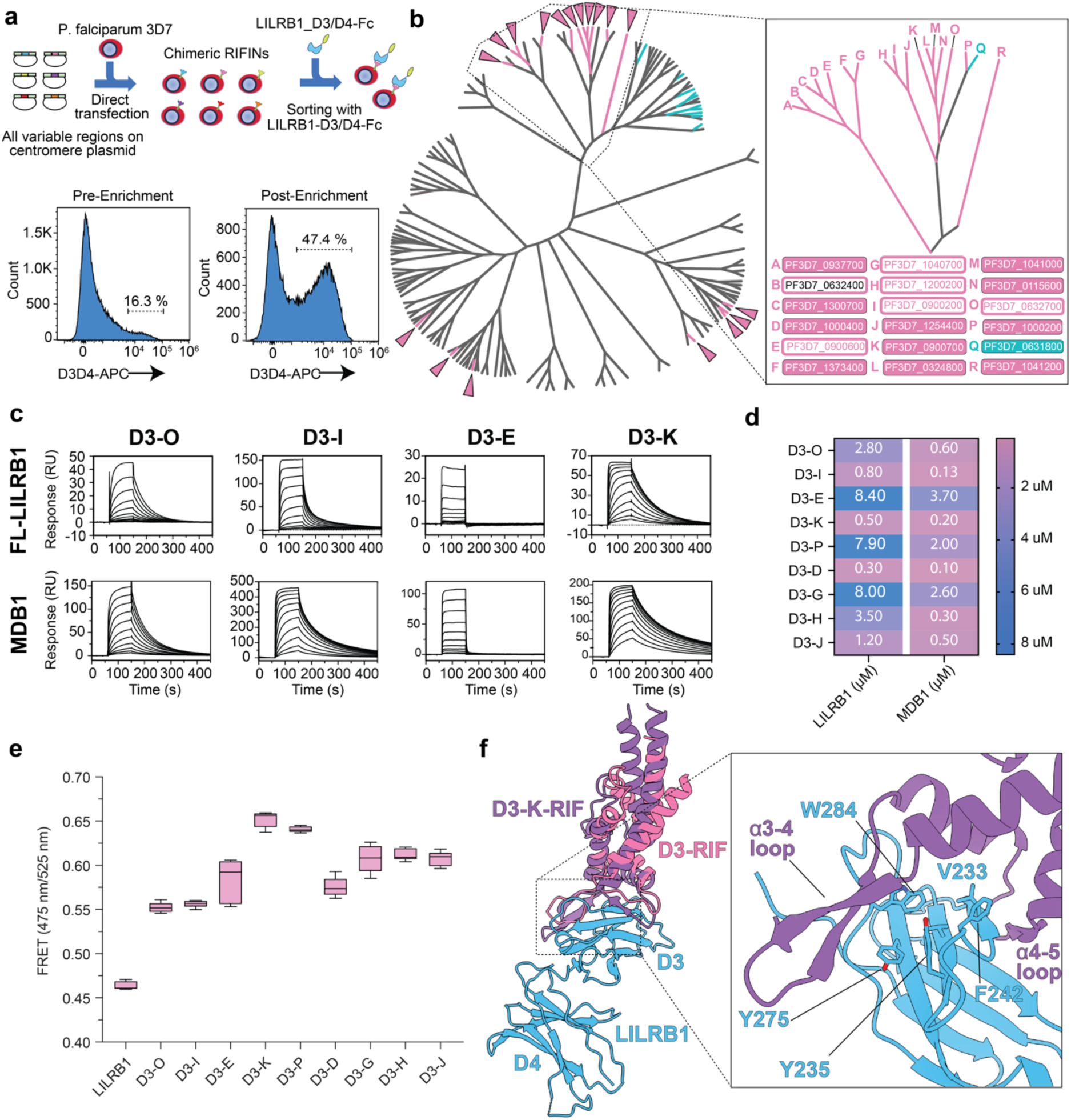
Identification of a clade of RIFINs bind to LILRB1 D3D4. **a)** The upper panel shows the generation of a library of *Plasmodium falciparum*-infected erythrocytes transfected to express individual RIFINs. These were selected by binding to LILRB1-D3D4-Fc. The lower panel shows the results of selection from pre-enrichment (left) to post-enrichment (right)**.a)** Phylogenetic tree of the entire repertoire of RIFINs from the 3D7 genome. Hits from two biologically independent library screening experiments are denoted with pink triangles and edges. KIR binding RIFINs previously described^11^ are shown with blue edges. The major LILRB1-D3D4 binding clade is shown in the inset (right) with a key for each D3-RIF (A-R) which provides the full names of the RIFIN. RIFINs in block colour boxes were identified in the library screening approach. Hollow boxes denote RIFINs confirmed to bind to LILRB1 by SPR. **c)** Representative SPR sensorgrams from multicycle experiments in which RIFINs were flowed over either LILRB1 (top) or MDB1 antibody (bottom). **d)** Summary of K_d_ values obtained from equilibrium fits of the data for all 9 RIFINs tested (n = 3, fits shown in Supplementary Figure 3e). **e)** FRET ratio 475 nm/525 nm of the emission spectrum of LILRB1 FRET probe after excitation at 433 nm either alone or in the presence of 10-fold molar excess of each RIFIN. Data from represent measurements from five independent experiments and two biologically independent preparations of the protein. Box and whisker plots show range, interquartile range and mean. **f)** Crystal structure of D3-K-RIF (purple) in complex with D3D4 of LILRB1 (blue) with D3-RIF (pink) overlayed to demonstrate the overall similarity of the complexes.

**Figure 3:**
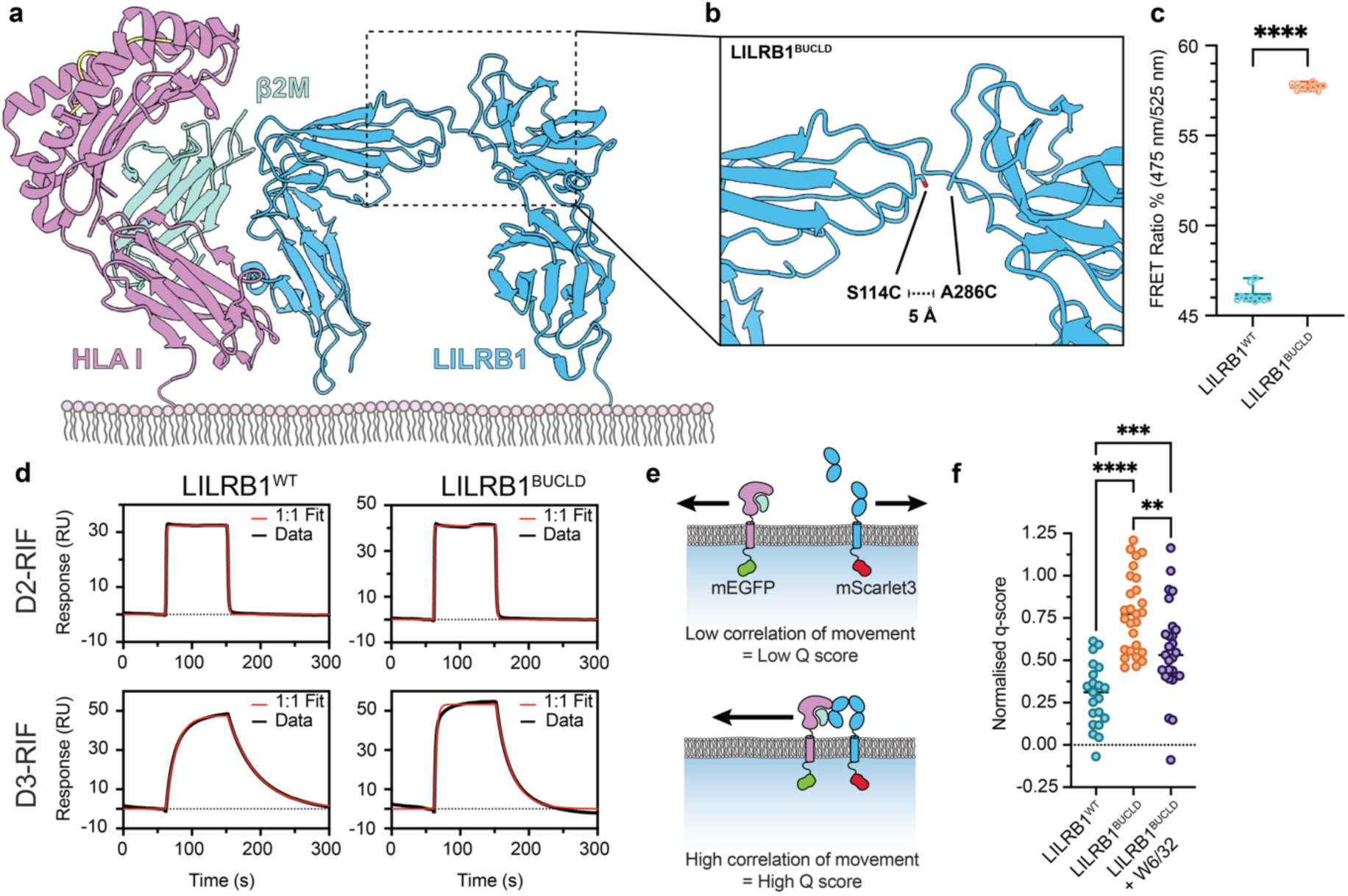
Buckled LILRB1 complex can form a complex with MHC class I on the same cell. **a)** Composite model showing how the LILRB1 buckled conformation revealed in this paper is compatible with a cis interaction with MHC class I, generated by taking superimposing the buckled LILRB1 structure onto a structure of LILRB1-D1D2 bound to MHC class I (PDB code 6EWA). **b)** A close up showing the two residues mutated to cysteine to form a disulphide bond in the LILRB1^BUCLD^ mutant and the distances between the hydroxyl group of Ser114 and the C_β_ of A286 in our structure. **c)** FRET ratio (475 nm/525 nm) of either LILRB1^WT^-FRET probe or LILRB1^BUCLD^-FRET probe after excitation at 433 nm. Datapoints represent measurements from ten independent experiments from at least two biologically independent preparations of the protein. Statistical significance was determined by one-way ANOVA in PRISM10. **** <0.0001. **d)** SPR sensorgrams (black) and kinetic fit to the data (red) for LILRB1^WT^ or LILRB1^BUCLD^ (analytes) bound to D2-RIF (ligand, top) or D3-RIF (bottom), representative of n = 3. **e)** Schematic of FCCS, where molecules moving separately have low *q* score, and molecules moving together have a high *q* score. **f)** FCCS measurements from HEK β2M^−/-^ transfected with HLA-A2-9V SCT mScarlet3 and either LILRB1^WT^-GFP or LILRB1^BUCLD^-GFP. Data show mean from three independent experiments, with data points indicating individual cells. LILRB1^WT^: 21 cells; LILRB1^BUCLD^: 27 cells; LILRB1^BUCLD^: 27 cells. Statistical significance was determined using an unpaired t-test. ****p < 0.0001.

To validate hits from library screening, transgenic iRBC lines were generated which express each of the identified LILRB1-D3D4 binding RIFINs from the major clade. Flow sorting of these lines showed that all bound to LILRB1-D3D4-Fc (Supplementary Figure 2c). To confirm that all members of the clade bind LILRB1-D3D4, we produced 10 more RIFINs from this clade, including five which were not hits in our library screen (*Pf*3D7_0632700, *Pf*3D7_1200200, *Pf*3D7_0900200, *Pf*3D7_0900600 and *Pf*3D7_1040700). We could express 9 of these RIFINs and used SPR to test their binding to MDB1 and LILRB1 (Figure 2c and Supplementary Figure 2d). All bound to both ligands, with higher affinities for MDB1 than for LILRB1 (Figure 2d and Supplementary Figure 2e).

To determine whether binding of all D3D4-binding RIFINs requires LILRB1 to adopt a buckled structure, we tested the panel against the LILRB1 FRET-sensor probe. All nine RIFINs increased the FRET ratio, showing that they increase LILRB1 buckling (Figure 2e). To determine whether differences in FRET ratio were due to different binding poses on LILRB1, the RIFIN with the highest FRET ratio (D3-K-RIF, *Pf*3D7_0900700) was crystallized in complex with D3D4 of LILRB1, allowing structure determination (Figure 2f). D3-RIF and D3-K-RIF occupy the same site on LILRB1 with the same loop between ⍺4 and ⍺5 forming an intermolecular β-sheet with β4 of the D3 domain of LILRB1 (Fig. 2F, Supplementary Figure 4). In both cases, the long loop between ⍺3 and ⍺4 of D3-K-RIF occupies the same hydrophobic pocket (Fig. 2f) which would otherwise mediate D2-D3 interdomain contacts and cannot therefore bind extended LILRB1. However, one difference is that the long binding loop of D3-K-RIF clashes with D2 of LILRB1 when LILRB1 is in the D3-RIF-bound conformation (Supplementary Figure 4b,c). This may cause the different FRET ratio between D3-RIF and D3-K-RIF, with these two RIFINs stabilising subtly different buckled conformations of LILRB1. Therefore, all LILRB1 D3D4-binding RIFINs are likely to target the same region of the D3 domain, but with subtle differences in the LILRB1 buckled conformation.

We next asked whether we can identify D3D4-binding RIFINs in a parasite genome isolated from Kenya. A phylogenetic tree comprising all RIFINs from both *Pf*3D7 and *Pf*KE01 strains was constructed (Supplementary Figure 5a) and six *Pf*KE01-derived RIFINs in the D3D4 clade were selected for recombinant expression. Five could be expressed in quantities sufficient for SPR and four of these bound LILRB1 and MDB1 (Supplementary Figure 5b). Therefore, phylogenetic analysis could be used to identify LILRB1-D3D4-binding RIFINs, albeit imperfectly. We also show that these D3D4-binding RIFINs are common not only in the 3D7 genome, but in genomes of a Kenyan strain, making it likely that they are widespread across strains of *Plasmodium falciparum*.

### A buckled conformation of LILRB1 interacts with MHC class I in cis

We next asked whether the buckled conformation of LILRB1 might be functionally relevant in NK cell signalling. Superimposing a crystal structure of LILRB1 D1D2 in complex with MHC class I ^20^ onto our buckled structure suggested that MHC class I and buckled LILRB1 can interact when both are presented from the same membrane, potentially allowing formation of a *cis* signalling complex (Figure 3a). To test this, we designed a form of LILRB1 locked into the buckled conformation (LILRB1^BUCLD^) by introducing a disulphide bond that bridges the D2 and D3 domains (Fig. 3b). LILRB1^BUCLD^ was assessed using size exclusion chromatography, revealing a monodisperse peak (Supplementary Figure 6a). The LILRB1^BUCLD^ mutation was tested in the context of the LILRB1 FRET probe, giving an increased FRET ratio when compared to LILRB1^WT^, showing successful stabilisation of a buckled conformation (Figure 3c). LILRB1^WT^ and LILRB1^BUCLD^ were each immobilized (∼300 RU) on a CM5 SPR sensor and D2 and D3 RIFINs were flowed across (Figure 3d). The binding of D2 RIF was unaffected by the presence of the mutation, demonstrating that the D1D2 interface is unaffected by locking. In contrast, fitting to a kinetic 1:1 model showed a ∼6-fold increase in the association rate (K_a_) for LILRB1^BUCLD^ compared to LILRB1^WT^ (2.82 × 10^5^ and 4.40 × 10^4^ 1/Ms, respectively). We attributed this rate increase to a requirement for LILRB1^WT^ to buckle before D3-RIF can bind, whereas the LILRB1^BUCLD^ mutant was constitutively buckled and poised to interact with the RIFIN more rapidly. Taken together these data demonstrate successful generation of a LILRB1^BUCLD^ mutant.

To investigate whether buckling of LILRB1 allows formation of *cis* interactions with MHC class I in a cellular context, we performed Fluorescence Cross-Correlation Spectroscopy (FCCS) experiments^21^. HEK cells lacking β2M were transfected with a single chain trimer form of MHC class I tagged with mScarlet3 at the C-terminus ^22^ and either LILRB1^WT^ or LILRB1^BUCLD^ tagged with mEGFP at the C-terminus. Areas of basal membrane were imaged to identify regions with uniform distributions of both proteins that are ideal for FCCS analysis (Figure 3e and f, Supplementary Figure 6b,c).

These regions were then subjected to line-scan FCCS, which provides the optimal balance of time resolution and statistical power for assessment of co-diffusion ^23^. We used a membrane anchored tandem fluorophore construct to determine the cross-correlation value (q) for a covalent dimer and set this q value to 100%, with the q value of non-interacting fluorophore constructs set to 0%. The normalised q score for LILRB1^WT^ was 0.29, indicating that around 30% of LILRB1^WT^ co-diffused with MHC Class I in a *cis*-complex on the cell surface. The normalised q-score increased to almost 0.75 for LILRB1^BUCLD^, indicating a greater proportion co-diffusing with MHC class I. Addition of the anti-MHC W6/32 mAb, which blocks the interaction between LILRB1^BUCLD^ and MHC class I, significantly diminished the normalized q-score to 0.55, suggesting that co-mobility is due to a specific interaction. These experiments show that LILRB1^BUCLD^ has a greater propensity to form *cis* interactions with MHC class I than LILRB1^WT^, although LILRB1^WT^ still interacts with MHC class I in *cis*, most likely when in the buckled conformation.

### The cis LILRB1-MHC Class I complex produces an inhibitory signal in immune cells

Immune receptor mediated signalling can be either *trans*, with the ligand on a different cell to the receptor or *cis*, with both ligand and receptor on the same cell. *Trans* signalling is often a direct process of active signalling. However, *cis* signalling is less well understood and has been proposed to set the basal level of cellular responsiveness to *trans* signalling ^24,25^. In some cases, the *cis*-interaction has been proposed not to signal, but instead to reduce the pool of receptors available to receive a *trans* signal through competition ^26^. In other cases, *cis* signalling has been proposed to provide low-level basal signalling that increases the threshold of positive signal required for cell activation ^27,28^. However, studying the functional effects of *cis* protein complexes has proved technically challenging due to difficulties in distinguishing *cis* and *trans* signalling, and functional *cis*-signalling has not been experimentally demonstrated ^29^.

Having revealed that LILRB1^BUCLD^ enhances the interaction of LILRB1 with MHC in *cis*, we asked if this also causes inhibitory signalling in NK cells. We first used a supported lipid bilayer (SLB) system to image immune synapses formed with the NK cell line, NK92 expressing either LILRB1^WT^ or LILRB1^BUCLD^ (Figures 4a,b). NK92 cells lack Fcγ receptors that would be used during antibody dependent cellular cytotoxicity of iRBC, but express NKp30, an activating receptor that shares signalling subunits with Fc receptors. To mimic contacts formed during cell-mediated cytotoxicity, we coupled ICAM-1, CD58 and B7-H6 to SLBs. ICAM-1 and CD58 form an adhesive interaction with LFA-1 and CD2, respectively, whilst B7-H6 engages NKp30 to activate the NK92 cells ^30^. NK92 cells expressing LILRB1^WT^ formed immune synapses and perforin was deposited at the synapse, indicating NK cell activation. However, for NK92 cells expressing LILRB1^BUCLD^, perforin deposition was lower, despite similar levels of LILRB1 expression (Supplementary Figure 7), suggesting inhibitory signalling mediated by buckled LILRB1. The W6/32 antibody, which blocks MHC class I from binding by LILRB1, led to increased perforin deposition in NK92 cells expressing LILRB1^BUCLD^ (Figure 4b). This suggests that buckled LILRB1 mediated inhibitory signalling and that this requires an interaction with MHC class I.

**Figure 4:**
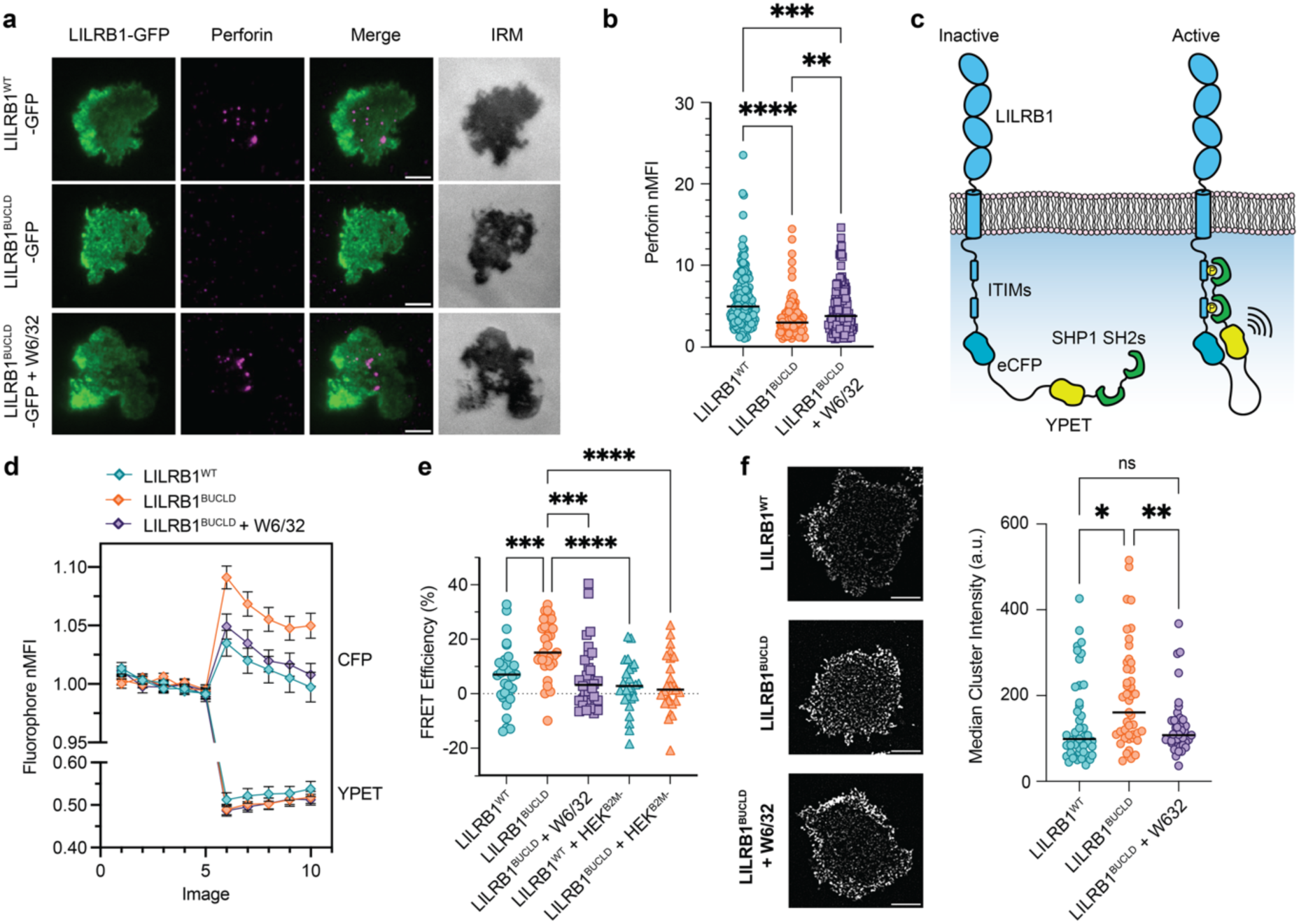
The LILRB1-MHC class I *cis* complex produces functional inhibitory signals in NK cells. **a)** Representative images and **b)** Analysed perforin secretion from the contact area of NK92 cells expressing LILRB1-GFP constructs, when incubated with supported lipid bilayers coated with ICAM1, B7-H6 and CD58 for 30 mins. Scale bars 5 µm. Data show the mean, with points indicating individual cells from 4 independent repeats. LILRB1^WT^: 147 cells; LILRB1^BUCLD^: 136 cells; LILRB1^BUCLD^ + W6/32: 140 cells. **c)** Schematic of FRET signalling reporter transfected into HEK293 cells used for **d)** and **e)**. **d)** Normalised fluorophore fluorescence intensity of photobleached region of interest from HEK293 cells transiently expressing either LILRB1^WT^- or LILRB1^BUCLD^ -containing FRET signalling reporter, before or after photobleaching. Data show mean ± SEM from three independent experiments. **e)** calculated FRET efficiency from **d)**. In both, data show median, with data points showing individual cells from 3 independent experiments. LILRB1^WT^: 31 cells; LILRB1^BUCLD^: 35 cells; LILRB1^BUCLD^ + W6/32: 32 cells; LILRB1^WT^ + HEK^B2M-^: 26 cells; LILRB1^BUCLD^ + HEK^B2M-^: 26 cells. **f)** Representative SIM images of NK92 basal membrane and analysis of LILRB1 cluster integrated density for the contact area of NK92 cells expressing either LILRB1^WT^, LILRB1^BUCLD^, or LILRB1^BUCLD^ with 10 µg mL^−1^ W6/32 antibody; when added to supported lipid bilayers coated with ICAM1, B7-H6 and CD58 for 30 mins. Data show mean, with datapoints indicating individual cells from 3 independent repeats. LILRB1^WT^: 45 cells; LILRB1^BUCLD^: 45 cells; LILRB1^Bucld^ + W6/32: 39 cells. Statistical significance was determined by one-way ANOVA with Tukey’s post-hoc multiple comparison test. *p < 0.05, **p < 0.01, ***p < 0.001, ****p < 0.0001.

In addition to studying the end point of perforin secretion, we set up a more direct assay to measure the phosphorylation status of the LILRB1 ITIMs as a read-out of receptor activation ^31^. We produced a reporter which contains a FRET module immediately after the ITIMs of LILRB1, consisting of ECFP, a spacer region, YPET and then the two SH2 domains of SHP-1. ITIM phosphorylation and subsequent SH2 engagement will bring the two fluorescent proteins into proximity, generating a FRET signal (Figure 4c, Supplementary Figure 8). HEK293 cells were transfected with forms of the FRET probe containing either LILRB1^WT^ or LILRB1^BUCLD^ together with the Lck kinase to mediate ITIM phosphorylation. To assess the degree of FRET signalling, accepter photobleaching was used to destroy the acceptor fluorophore and the divergence in donor and acceptor fluorescence was used assess the degree of FRET (Figures 4d,e and Supplementary Figure 8). The FRET signal for LILRB1^BUCLD^ was greater than that for LILRB1^WT^, indicating greater ITIM phosphorylation. Indeed, repeating this assay for LILRB1^BUCLD^ in the presence of W6/32 antibody or for both LILRB1^BUCLD^ and LILRB1^WT^ in HEK β2M^−/-^ cells, all of which disrupt the interaction of LILRB1 with MHC class I, resulted in a similar signal to LILRB1^WT^ (Figure 4e). Taken together, these data show the interaction of LILRB1 with MHC class I in *cis* provides an inhibitory signal.

Immune signalling is often driven by cluster formation at immune synapses. However previous experiments using the FRET Phosphosensor were conducted by imaging areas of membrane which lack visible clusters. We therefore imaged NK92 cells expressing either LILRB1^BUCLD^ or LILRB1^WT^ using Structured Illumination Microscopy (Fig. 4f). We observed more LILRB1 dense clusters of LILRB1^BUCLD^ than of LILRB1^WT^ and few LILRB1^BUCLD^ clusters were observed in the presence of W6/32. These data suggest that MHC class I induces LILRB1 clusters through formation of a *cis* complex.

These data show that LILRB1 can form complexes with MHC class I in *cis* which cause ITIM phosphorylation and microcluster formation and which suppress perforin secretion. In each case, these are increased by locking LILRB1 in a buckled conformation and are inhibited by the blocking W6/32 antibody. These data indicated formation of a complex of LILRB1 with MHC class I in *cis*, which inhibits immune cell activation.

### D3-binding RIFINs can signal through a complex of LILRB1 and MHC class I

As we find that buckled LILRB1 can mediate inhibitory signalling through a *cis* interaction with MHC class I and that D3-binding RIFINs can signal through LILRB1, we next tested whether D3-binding RIFINs can signal through buckled LILRB1 bound to MHC class I. We first performed a bilayer interferometry (BLI) assay to determine whether D3-binding RIFINs and MHC class I can simultaneously bind to LILRB1 (Figure 5a). Equal densities of biotinylated LILRB1 were immobilised on streptavidin sensors, either alone, or in the presence of two different concentrations of biotinylated MHC class I. Biotin placement allowed immobilisation of LILRB1 and MHC class I on the same sensor to mimic cell-surface presentation and allow *cis* complex formation. Sensors were then exposed to 10 µM D2-RIF or D3-RIF. Increasing the amount of immobilized MHC class I progressively reduced binding of D2-RIF, as MHC class I and D2-RIF bind overlapping sites on LILRB1. In contrast, binding of D3-RIF increased with more immobilised MHC class I, most likely due to MHC class I-mediated stabilisation of buckled LILRB1 (Figure 5a). Indeed, when repeated using LILRB1^BUCLD^, D3-RIF binding was unaffected by MHC class I density (Figure 5a). These data demonstrate that *cis* MHC class I can either inhibit or promote LILRB1 from binding to RIFINs, depending on whether the RIFIN binds D2 or D3.

**Figure 5:**
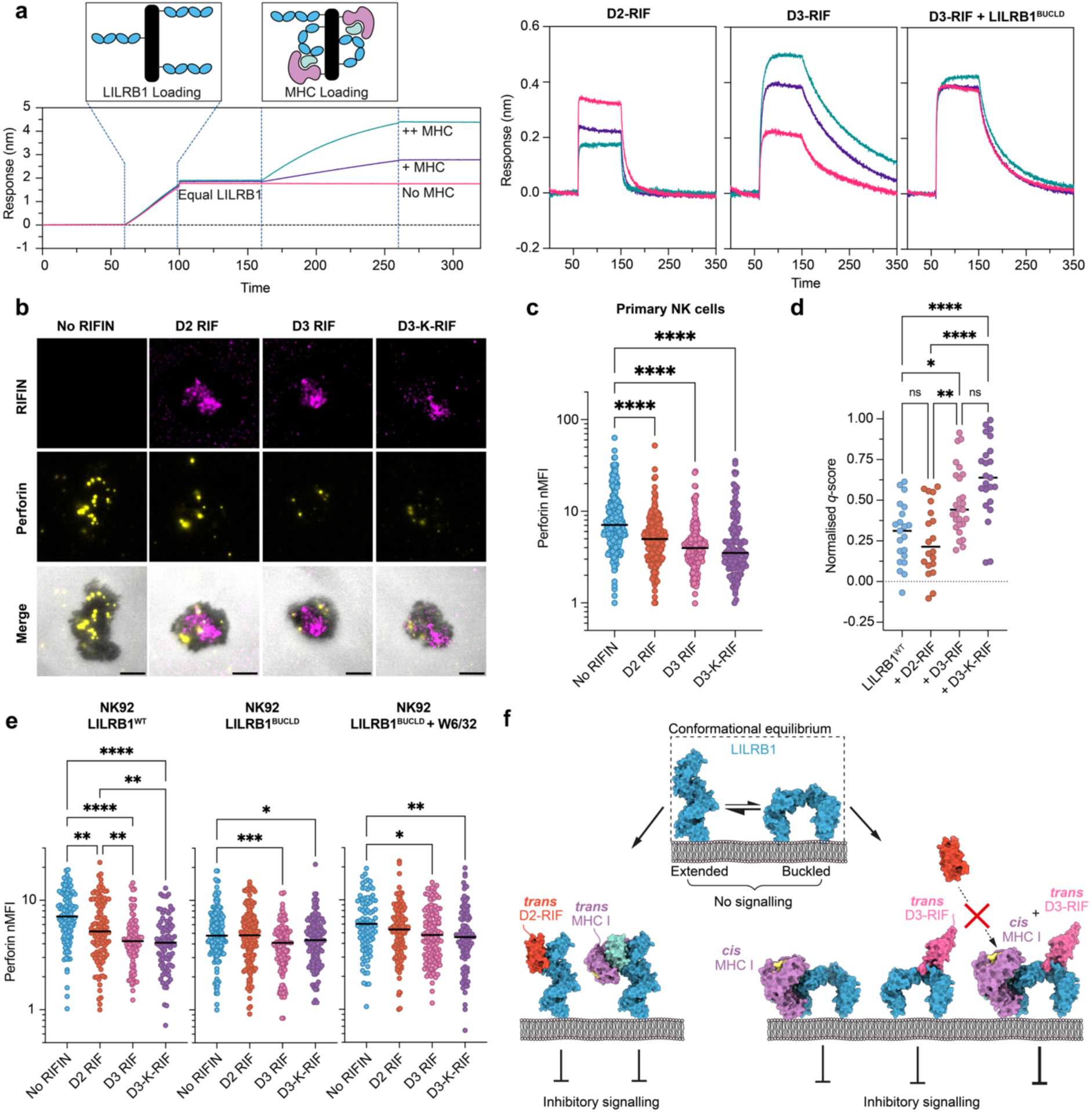
D3-RIF but not D2-RIF can signal through a cis LIRB1-MHC class I complex. **a)** The left-hand panel shows BLI response curves for loading of biotinylated LILRB1 and MHC class I onto three SA sensors. The biotinylation site is at the site of normal membrane attachment to mimic native presentation of the ligands as a *cis* complex. The right-hand panel shows injection of D2- or D3-RIF over sensors with immobilised LILRB1^WT^ and varying amounts of MHC class I (left and centre) and D3-RIF over sensors or with immobilised LILRB1^BUCLD^ and varying amounts of MHC class I (right), each representative of n=3. **b)** Representative images and **c)** Analysed perforin secretion from the contact area of NK cells when added to activating supported lipid bilayers coated with D2-RIF, D3-RIF, D3-K-RIF or no RIFIN for 30 mins. Scale bars 5 µm. Data show median, with data points indicating individual cells from 4-7 independent donors. No RIFIN: 239 cells; D2-RIF: 183 Cells, D3-RIF: 193 cells; D3-K-RIF: 134 cells. **d)** FCCS measurements from HEK β2M^−/-^ transfected with HLA-A2-9V SCT mCherry2 and LILRB1^WT^-GFP in the presence of the specified antibody or RIFIN. Data show mean from three independent experiments, with data points indicating individual cells. LILRB1^WT^: 21 cells; LILRB1^WT^ + D-RIF: 20 cells; LILRB1^WT^ + D3-RIF: 27 cells; LILRB1^WT^ + D3-K-RIF: 23 cells. **e)** Analysed perforin secretion from the contact area of NK92 cells expressing either LILRB1^WT^-GFP (left) LILRB1^BUCLD^-GFP (centre), or LILRB1^BUCLD^ GFP (right) with 10 µg mL^−1^ W6/32 antibody (right); when added to supported lipid bilayers coated with ICAM1, B7-H6 and CD58 and 100 molecules µm^−2^ of the specified RIFIN for 30 mins. Scale bars 5 µm. Data show mean, with data points indicating individual cells from 4 independent repeats. LILRB1^WT^: 113 cells; + D-RIF: 113 cells; + D3-RIF: 97 cells; +D3-K-RIF: 90 cells. LILRB1^BUCLD^: 152 cells; + D2-RIF: 148 cells; + D3-RIF: 133 cells; + D3-K-RIF: 136 cells. LILRB1^BUCLD^ + W6/32: 89 cells; + D2- RIF: 132 cells; + D3-RIF: 135 cells; + D3-K-RIF: 117 cells. Statistical significance was determined by one-way ANOVA with Tukey’s post-hoc multiple comparison test. *p < 0.05, **p < 0.01, ***p < 0.001, ****p < 0.0001. **f)** Schematic summary of the findings of this paper.

We next used the SLB assay to test whether D3-binding RIFINs can signal through LILRB1 bound to MHC class I. We first tested whether D3-binding RIFINs can suppress perforin release from primary NK cells using our SLB assay (Figures 5b,c and Supplementary Figure 9). We incubated NK cells with SLBs containing IgG and ICAM1 to induce immune synapse formation, with the Fc region of the IgG engaged by CD16 on the NK cells. These bilayers contained D2-RIF, D3-RIF, D3-K-RIF or no RIFIN. All three RIFINs reduced perforin deposition, with D3-RIF and D3-RIF-K trending towards lower perforin deposition, with median nMFI of 3.96 and 3.51, respectively, compared to 4.98 for D2 Rif. Therefore D3-binding RIFINs can suppress activation of primary NK cells.

As D3-binding RIFINs stabilise the buckled form of LILRB1 in solution, we also tested whether they increase the association of LILRB1 with MHC class I in *cis*. We used HEK293 cells lacking β2M and expressing a single chain HLA fused to mScarlet3 and LILRB1-GFP to conduct FCCS measurements to assess their degree of co-localisation (Figure 5d). Whilst D2-RIF did not increase cross-correlation of LILRB1 and MHC I compared to no RIFIN, both D3-RIF and D3-RIF-K caused a significant increase in cross-correlation, with the average q-score increasing from 0.29 for LILRB1^WT^ to 0.49 and 0.64 for D3-RIF and D3-RIF-K respectively. Therefore D3-binding RIFINs increase the amount of buckled LILRB1 and its binding to MHC class I in *cis*, thereby increasing inhibitory signalling through this complex.

Finally, we tested the effect of LILRB1 buckling on signalling due to D3-binding RIFINs. NK92 cells were transferred with either LILRB1^BUCLD^ or LILRB1^WT^ and these were incubated with SLBs containing D2-binding or D3-binding RIFINs (Figure 5e). For LILRB1^WT^, D2-RIF inhibited perforin deposition, reducing the median nMFI from 7.11 with no RIFIN to 5.2, however both D3-binding RIFINs were more potent inhibitors, with median nMFIs of 4.24 for D3-RIF and 4.10 for D3-K-RIF. In an equivalent experiment with LILRB1^BUCLD^, inhibition of perforin deposition was observed without any RIFIN and was not increased by D2-RIF. However, D3-RIF and D3-K-RIF both significantly increased inhibitory signalling by reducing perforin release. When the same experiment was conducted using LILRB1^BUCLD^ in the presence of W6/32, D3-RIF and D3-K-RIF were less inhibitory, showing that the interaction between LILRB1^BUCLD^ and MHC class I enhances, but is not required for inhibitory signalling in NK cells.

## Discussion

The ancient malaria parasite, *Plasmodium falciparum*, has evolved in the context of the human immune system. Despite replicating within human erythrocytes, it remains susceptible to destruction by immune cells, such as NK cells and macrophages ^5,6,32–34^. As a result, it evolved a large family of RIFIN proteins which it deploys to infected erythrocyte surfaces. RIFINs have been identified which bind to inhibitory immune receptors LILRB1^8,17^, LILRB2^9^, LAIR1^10,35,36^ and KIRs^11^ and which signal through these receptors to suppress NK cell activation and perforin deposition^11,17^. In this study, we focused on RIFINs which bind to LILRB1 and resolved a mystery. While some RIFINs bind to the D2 domain of LILRB1 and trigger inhibitory signalling, others bind to the D3 and D4 domains of LILRB1 which have been inserted into antibodies^12^. Here we confirm that there are indeed two separate clades of LILRB1 binding RIFINs. Furthermore, we show that the D2-binding RIFINs are compatible with an elongated conformation of LILRB1 while LILRB1 must buckle to bind to D3-binding RIFINs. These two distinct groups of RIFINs are both able to signal through LILRB1 and reduce NK cell activation.

This raised the question of why a group of RIFINs has evolved to bind to the buckled conformation of LILRB1 when only elongated LILRB1 has previously been observed. Is the buckled conformation of LILRB1 found on immune cells and does it signal? We show here that indeed the buckled conformation of LILRB1 is present on NK cells, allowing interaction between LILRB1 and MHC class I molecules on the same cell surface. We also show that these complexes provide a low-level inhibitory signal in ‘*cis*’ to NK cells. This *cis* signal will reduce the basal activity of NK cells and increase the threshold which must be overcome by *trans* signals to trigger perforin deployment. LILRB1 therefore exists in a dynamic equilibrium on NK cells, in either elongated or buckled conformations (Figure 5f). Increased concentrations of MHC class I in *cis* will increase the amount of buckled LILRB1 and increase the amount of basal inhibitory ‘*cis*’ signalling. Ligands *in trans*, including MHC class I on a target cell, will favour the elongated conformation of LILRB1, generating increased signalling in ‘*trans*’. This allows the immune system to tune the activity of LILRB1 and the susceptibility of immune cells to activation by using ligands in ‘*cis*’ or ‘*trans*’ to regulate the conformation of LILRB1 and to set the responsiveness of NK cells. This is the first time that *cis*-signalling through an inhibitory receptor has been experimentally demonstrated to reduce immune cell function but it has the potential to be a common mechanism.

Together, these data show that two clades of RIFINs have evolved which interact with one or other of the naturally occurring conformations of LILRB1. In each case, the RIFINs can induce inhibitory signalling when provided in ‘*trans*’, either through elongated LILRB1, or through buckled LILRB1 in complex with MHC class I (Figure 5f). Each clade of RIFIN is thereby able to achieve the same objective of suppressing NK cell function, whether the quantity of MHC class I on that immune cell favours elongated or buckled LILRB1. This study gives another example of the breadth of the RIFIN repertoire to function through different inhibitory receptors in different ways, and their study has revealed a novel inhibitory mechanism mediated by LILRB1. With around half of the RIFINs still uncharacterised, more RIFIN receptors and mechanisms of immune suppression likely remain to be discovered.

## Materials and Methods

### Protein Expression and Purification

All RIFIN genes were synthesized (ThermoFisher, Geneart) and subcloned into either pHLsec, containing a C-terminal His_6_ tag (D3-RIF, residues 173-326, and D3-K-RIF, residues 178-331 G185C T317C) for structural and functional experiments or pENTR4LP for D3-K-RIF production and SPR screening. All proteins were harvested 5 days after transfection of Expi293F™ GnTI- cells with the ExpiFectamine™ transfection kit (ThermoFisher Scientific). Supernatants were harvested by pelleting the cells at 5,000 xg before filtering with a 0.45 µm bottle-top filter. For His-tagged proteins, filtered supernatant was passed over Ni Sepharose® Excel resin (Cytiva) before washing with 50 mM Tris 8.0, 150 mM NaCl, 20 mM imidazole and eluting with 50 mM Tris 8.0, 150 mM NaCl, 500 mM imidazole. Eluted protein was concentrated and digested with EndoHF overnight before application to a Superdex S75 10-300GL column (Cytiva) for final purification. RIFINs expressed in the pENTR4LP vector were cloned between an N-terminal, cleavable, monomeric Fc and a C-terminal C-tag according to the domain boundaries in Table S2. Cells were transfected as above and the filtered supernatant added to CaptureSelect C-tagXL Affinity Matrix (Thermo Fisher Scientific) for affinity purification. Resin was washed with HBS (20 mM HEPES, 150 mM NaCl pH 7.5) before elution with 2M MgCl_2_, 20 mM Tris 7.5. Eluted protein was concentrated and buffer exchanged back into HBS before overnight TEV and EndoH_f_ cleavage to simultaneously remove both monomeric Fc and glycans. After cleavage, C-tagXL resin was used to remove cleaved monoFc before once more eluting the pure protein and polishing was performed using size exclusion chromatography.

All LILRB1 variants used, including FRET-sensors and D3D4 LILRB1 were subcloned into pHLsec vector. For Avi-tagged variants the vector was modified to replace the usual C-terminal His_6_-tag with an Avitag and the His_6_-tag was placed at the N-terminus after the secretion signal. Constructs were expressed in HEK293F cells transfected using polyethyleneimine and purified as above (without EndoHF treatment). LILRB1-D3D4-Fc for flow sorting experiments was expressed in adherent HEK293T cells (Riken BRC, RCB2202) using TransIT-293 (MirusBio) and the cell supernatant used for experiments.

HLA Cw4 was expressed and purified as previously described using a refolding approach from bacterial inclusion bodies ^11^.

### Crystallization, data collection and structure determination

FL-LILRB1 and D3-RIF were combined at a 1:1 molar ratio and purified using size exclusion chromatography (Superdex S75 10/100GL, Cytiva). Initial hits grew between 49 and 56 days with a well solution containing 2.8 M sodium acetate trihydrate pH 7.0. Crystals were further optimized through an additive screen which included seeds from the original crystals. The best diffracting crystals were obtained with 100 nL of well solution (2.8 M sodium acetate trihydrate pH 7.0), 100 nL of complex (10 mg/mL), 25 nL of Silver Bullets condition H7 (Hamptons Research) and 25 nL of seeds. Crystals were cryoprotected in 4M sodium acetate + 20% glycerol before freezing in liquid nitrogen.

Data were recorded on the i04 beamline at Diamond Light Source at a wavelength of 0.9795 Å and processed using the automatic Xia2 3dii pipeline. The structure was solved by molecular replacement using D1D2 and D3D4 di-domains of LILRB1 and the previously published RBL1D3 RIFIN ^52^ (PDB accession codes 6ZDX and 7KFK, respectively) as search models, using Phaser MR (v.2.8.3) ^46^. Building and refinement cycles were carried out using COOT (v.0.8.9.2)^48^ and PHENIX (v.1.2.12)^53^.

For the complex of D3-K-RIF and LILRB1-D3D4, EndoHF treated D3-K-RIF and LILRB1-D3D4 (residues 178 – 331) were combined and eluted as a 1:1 complex from a size exclusion column in 20 mM HEPES pH 7.5, 150 mM NaCl. Initial crystals formed at 10 mg/mL in F1 of the JCSG Core I screen (Molecular dimensions). Fine screening around this condition led to diffraction quality crystals at 10 mg/mL protein complex in 100 mM MES, pH 5.2, 10% PEG 6000 + B9 of the Silver bullet screen (Hamptons Research). Crystals were flash frozen in the reservoir solution (no silver bullet) + 20% glycerol. Data were recorded on the i04 beamline at Diamond Light Source at a wavelength of 0.9537 Å. Data were truncated to 2.8 Å using AIMLESS and molecular replacement was carried out using LILRB1-D3D4 (PDB 6ZDX) and an AlphaFold 3 predicted structure of D3-K-RIF ^50^. Building and refinement cycles were carried out using COOT (v.0.8.9.2)^48^ and BUSTER (v.2.10)^54,55^.

### Surface Plasmon Resonance

All SPR experiments were carried out in 20 mM HEPES pH 7.5, 300 mM NaCl, 0.005% v/v TWEEN-20 with a Biacore T200 instrument and evaluated in BIAevaluation software (Cytiva). For experiments to test the function of LILRB1 locking mutations, ∼700 RU of each LILRB1 variant was immobilized to a flow cell of a CM5 sensor after activation using EDC/NHS. D3-and D2-RIFs were injected across the sensor surface in a multicycle experiment spanning a concentration range of 60 nM to 4 µM. For screening of 3D7 and Kenyan RIFINs, 600 RU of FL-LILRB1 was immobilized on Fc2 and 1200 RU of MDB1 on Fc3. Dilution series of each RIFIN were flowed across the sensor with starting concentrations ranging from 10 to 40 µM. In all cases, regeneration of the sensor surface was carried out using a 10 s pulse of 10 mM glycine pH 2.5 in-between cycles. Sensorgrams were double referenced with buffer blank and non-immobilized flow channel sensorgrams being subtracted from the raw data.

### FRET experiments

All FRET experiments were carried out using a CLARIOstar PLUS microplate reader (BMG Labtech), with 96 well non-treated black plates (ThermoFisher Scientific) with 200 µL per well in 20 mM HEPES 7.5, 150 mM NaCl. A LILRB1 FRET-sensor was designed using an existing molecular crowding sensor as a template^56^. Emission spectra of a concentration series from 1 µM to 4 nM was measured at 475 nm (mCerulean) and 525 nm (mCitrine) to ensure there was no auto-FRET in this concentration range. Subsequent experiments were carried out with the FRET sensor at 1 µM. The titration was carried out for concentrations of RBL1D3 from 16 µM to 31 nM, and the data was fitted to a 1:1 binding model in PRISM 9. For testing the ability of RIFINs to buckle the LILRB1 structure, RIFINs were added to the LILRB1 FRET sensor (0.5 µM) to a final concentration of 6 µM. Experiments with LILRB1 mutants were carried out again with the protein at 1 µM.

### Bilayer Interferometry

Bilayer interferometry (BLI) measurements were recorded on a OctetRed 384 instrument (FortéBio) using Octet SA biosensors (Sartorius). All sensorgrams were referenced by subtracting non-specific binding signal to an SA sensor with no protein immobilized on the surface. LILRB1-Avitag, LILRB1^BUCLD^-Avitag and HLA Cw4-Avitag were specifically biotinylated using the BirA500 kit (Avidity LLC) before buffer exchange into BLI buffer (20 mM HEPES 7.5, 150 mM NaCl, 0.005% v/v TWEEN-20) using a CentriPure P5 desalting column (Genaxxon Bioscience). Ligands were then sequentially immobilized to the SA surface, with equal quantities of LILRB1 (WT or BUCLD), followed by either no ligand, or increasing responses of HLA Cw4. Binding was tested for 60 s equilibration in buffer, followed by a 90 s association step in D2- or D3- RIFIN at 5 µM and a 300 s dissociation step. Experiments were performed in triplicate and referenced sensorgrams were plotted in PRISMv10.

### Transfection of parasites

*P. falciparum* culture and transfection were performed as previously described ^11^. Briefly, parasites were maintained in human O-type erythrocytes at 2% hematocrit in complete RPMI 1640-based medium (CM) supplemented with 10% human serum and 10% AlbuMAX I (Invitrogen), 25 mM HEPES, 0.225% sodium bicarbonate and 0.38 mM hypoxanthine under 90% N_2_, 5% CO_2_ and 5% O_2_. Parasites were regularly synchronized using sorbitol treatment.

For transfection, highly synchronized schizonts of the 3D7 strain were purified using a Percoll/sorbitol gradient solution and transfected with pFCEN-rif plasmids containing human dihydrofolate reductase gene as drug selectable marker. Transgenic parasites could be selected with pyrimethamine and used for each experiment. All transfections were performed in duplicate to obtain biologically independent lines.

### Generation and screening of RIFIN library

The generation of the RIFIN library was performed as described previously ^11^. Briefly, DNA fragments encoding all RIFIN variable regions were amplified from genomic DNA of the 3D7 strain using three degenerated primers (STAR methods) and were cloned into pFCEN-rif. The resulting plasmid pool was introduced into the 3D7 strain by electroporation. Transfected parasites were selected by treatment with pyrimethamine, generating a RIFIN library in which individual parasites expressed distinct RIFIN variants. Two biological independent libraries were produced and used for the study.

The iRBCs from each library were incubated with LILRB1-D3D4–Fc, followed by staining with a PE-conjugated anti-human IgG Fc antibody (Jackson ImmunoResearch, 109-116-170), and subjected to cell sorting (SONY, SH800). Sorted parasites were immediately returned to culture in CM and plasmid DNA was recovered. Inserted variable regions were amplified using primers targeting the vector backbone. To introduce sequencing adapters, a second PCR was performed using amplified variable regions as a template. The resulting sequence-tagged PCR products were sequenced on a MiSeq platform (Illumina).

Sequencing reads corresponding to RIFIN variable regions were extracted from FASTQ files using SeqKit with the ‘grep’ option, and vector-derived sequences were removed using Trimmomatic 0.39-2. After quality filtering, reads were aligned to the *P. falciparum* 3D7 reference genome (PlasmoDB v68) using Bowtie2 and reads mapping to multiple loci were excluded. The number of reads for each RIFIN were counted using featureCounts 2.0.1 and normalized to total mapped reads. Enrichment was determined relative to the pre-selection library, and RIFINs exhibiting greater than twofold enrichment were defined as candidate D3-RIFs.

Candidate D3-RIFs were amplified from 3D7 genomic DNA, individually cloned into pFCEN-rif, and introduced into parasites. Binding of each transgenic parasite to LILRB1 D3D4–Fc was assessed by flow cytometry using a CytoFLEX instrument (Beckman Coulter) with an APC-conjugated anti-human IgG Fc antibody (Jackson ImmunoResearch, 109-136-098).

### Assay for NKL cytotoxicity

The stable transfectant of K562 with D3-RIF (PF3D7_1373400) was generated using a retrovirus system as described previously ^11^. The inhibitory effect of D3-RIF on NK cell cytotoxicity was evaluated using NKL cells which constitutively express LILRB (provided by Dr. L. L. Lanier and Dr. H. Arase). Prior to the assay, viability of both NKL and K562–D3-RIF cells were assessed. If the viability of cells was below 85%, they were purified using Ficoll-Paque PLUS (Cytiva, 17144003) and cultured for an additional 3 days. Target K562–D3-RIF cells with high viability were labelled with 15 μM calcein acetoxymethyl ester (calcein-AM) in assay medium (phenol red–free RPMI 1640 supplemented with 1% FCS) for 30 min at 37 °C, followed by two washes. Effector NKL cells were similarly washed and co-incubated with target cells in 96-well plates at effector-to-target ratios ranging from 3:1 to 30:1. Plates were centrifuged at 250g for 5 min to promote cell contact and incubated for 4 h at 37 °C. Maximal calcein release was determined by lysing target cells with 1% Triton X-100 for 30 min before the end of incubation, while spontaneous release was measured from target cells cultured in the absence of NKL cells. Following co-incubation, plates were centrifuged at 1,500 rpm for 2 min, and fluorescence in the supernatant was quantified. Specific cytotoxicity C (%) was calculated using the following formula: C = 100 × (mean fluorescence in co-culture − mean fluorescence in spontaneous lysis)/(mean fluorescence in maximal lysis − mean fluorescence in spontaneous lysis). Parental K562 cells and K562-D2-RIF (PF3D7_1254800) cells were used as controls. All experiments were performed in quadruplicate.

### Phylogenetic analysis of RIFINs

Amino acid sequences for the variable regions of all 3D7 RIFINs were obtained from PlasmoDB (https://plasmodb.org/plasmo/app/) and alignment carried out using Clustal Omega (https://www.ebi.ac.uk/jdispatcher/msa/clustalo). Phylogenetic trees were produced with R using the ggtree module ^57^. For the prediction of LILRB1-D3D4-binding RIFINs from the Kenyan isolate *Pf*KE01, 3D7 RIFIN sequences confirmed to bind D3D4 using the library screening approach were combined with the RIFIN variable sequence repertoire from the *Pf*KE01 genome and same alignment and tree generation approach was used as above.

### Cell culture

NK92 cells (provided by Dr Luisa Burger) were cultured in alpha-MEM (αMEM, Gibco, 22571) supplemented with 12.5 % v/v foetal bovine serum (Gibco, A5456801), 12.5 % v/v equine serum (Gibco, 26050088), 2 mM L-Glutamine (Gibco, 25030) and 250 mg L^−1^ Gentamicin (Gibco, 15750). αMEM media was supplemented with 50 U mL^−1^ of human IL-2 (Peprotech, 200-02) upon culturing the cells. HEK 293 cells (ATCC, CRL-3216) were grown in complete Dulbecco’s Modified Eagle Medium (DMEM, Gibco, 11960), supplemented with 10% v/v FBS, 2 mM L-Glutamine, 1 mM Sodium Pyruvate (Gibco, 11360), 100 U mL^−1^ Penicillin (Gibco, 15140), 100 μg mL^−1^ Streptomycin (Gibco, 15140). Cells were cultured at 37 °C and 5 % CO_2_ in a humidified incubator.

### CRISPR-Cas9 RNP complex preparation

0.2 nmol LILRB1 crRNA (IDT, Hs.Cas9.LILRB1.AA) and 0.2 nmol Alt-R CRISPR-Cas9 tracrRNA (IDT, 1072543) were mixed in a volume of 2 µL and then sterilised by incubating at 95 °C for 5 mins. To this, 0.15 nmol of Alt-R S.p. Cas9 Nuclease V3 (IDT, 10000735) was added in 7.5 µL of nuclease-free Duplex Buffer (IDT, 1072570). This was incubated at 37 °C for 15 mins. 0.2 nmol of Alt-R Cas9 Electroporation Enhancer (IDT, 10007805) was then added and the RNP complex was electroporated into NK92 cells.

### NK92 electroporation

NK92 cells were electroporated using the Invitrogen Neon system. NK92 cells were pelleted at 150g for 7 mins. Cells were then resuspended in 5 mL αMEM without antibiotics and transferred to a 15 mL tube. This was centrifuged at 150g for 7 mins. Supernatant was then removed by pipette and the cells resuspended in Buffer R (Invitrogen) at 33.3×10^6^ cells mL^−1^. 3 mL of electrolytic buffer (Invitrogen, buffer E) was added to each Neon tube, which was then placed in the electroporation apparatus. For CRISPR/Cas9 KO of proteins; 9.5 µL of RNP complex was added to a 1.5 mL tube, followed by 120 µL of NK92 cells. For transgenic gene expression, 8 µg of DNA and 1 µg of PiggyBac mRNA (GenScript, RP-A00046) in a total volume less than 12 µL was added to a 1.5 mL tube, followed by 120 µL NK92 cells. The cells were then collected in a Neon tip on a Neon pipette, with the pipette then inserted into the Neon pipette station. The cells were then electroporated at 1650 V for 20 ms followed by 500 V for 100 ms. Cells were then pipetted into a 12 well plate, with 1 mL of αMEM without antibiotics added after 10 mins. This was incubated at 37 °C for 2 hours before 2 mL αMEM supplemented with 100 U mL^−1^ IL-2 was added to the cells before sorting by FACS.

### Preparation of HEK293 cells lacking β_2_ microglobulin

Lentivirus was produced by transfecting 1×10^6^ HEK293 cells in a 6 well plate with 1 µg pMDG, 1 µg p8.91 and 1 µg pLentiCRISPRv2_β2M_1 (GenScript, B2M CRISPR Guide RNA 1) using GeneJuice Transfection Reagent (Merck, 70967). Transfected cells were then incubated for 3 days to produce virus. 3 mL of virus-containing supernatant was then incubated with 1×10^6^ HEK293 cells in a 6 well plate. HEK293 β2-microglobulin knockout cells were sorted by FACS and confirmed by flow cytometry. p8.91 (Addgene 187441) and pMDG (Addgene 187440) were gifts from Simon Davis^58^.

### Isolation of Human NK cells

NK cells were isolated from anonymised leukocyte cones from healthy adult donors obtained through the NHS blood and transplant service under ethics agreement number 11-H0711-11, using RosetteSep Human NK Cell Enrichment Cocktail (StemCell Technologies, #15065). 50 µL of RosetteSep enrichment cocktail was added to 1 mL of leukocyte cone blood and incubated at RT for 20 mins. NK cells were enriched by density gradient centrifugation using Ficoll-Paque PLUS (Cytiva, 17144003), centrifuging at 1200g. NK cells were then washed once in PBS before red blood cell lysis using ACK lysis buffer (Gibco, A10492) as per the manufacturers’ instructions. NK cells were then washed twice with HEPES-buffered saline + 0.1 % w/v BSA + 1 mM CaCl_2_ + 2 mM MgCl_2_ (HBSS) before being resuspended at 5×10^6^ cells mL^−1^ in HBSS.

### Flow cytometry

Primary antibodies were anti-HLA-A,B,C Alexa Fluor 647 (Biolegend, 311414) and anti-LILRB1 (R&D systems, AF2017). 1×10^6^ cells were stained with LIVE/DEAD fixable aqua (Thermo Fisher, L34957) for 30 mins at 4 °C and then washed twice with PBS + 0.1 % w/v BSA. The cells were then stained with primary surface staining antibodies for 30 mins at 4 °C, prior to washing twice with PBS + 0.1 % w/v BSA. If required, cells were then stained with Alexa Fluor 647 AffiniPure F(ab) Fragment Donkey anti-Goat IgG (Jackson ImmunoResearch, 705-607-003) for 30 mins at 4 °C and then twice with PBS + 0.1 % w/v BSA. Cells were fixed in 4 % paraformaldehyde (PFA, ThermoFisher, 28908) v/v in PBS, before washing twice with PBS. Cells were then analysed on a Cytek 3-laser Northern Lights spectral flow cytometer using SpectroFlo v3.3 (Cytek). Further analysis was then performed with FlowJo v10.10.0 (BD Biosciences).

### Supported lipid bilayer assays

SLB experiments were performed as previously described ^17^. Briefly, 1,2-dioleoyl-*sn*-glycero-3-phosphocholine (Avanti Polar Lipids, 850375) micelles were supplemented with either 5 % or 2.5 % 1,2-dioleoyl-sn-glycero-3-[(N-(5-amino-1-carboxypentyl) iminodiacetic acid) succinyl]-Ni (Avanti Polar Lipids, 790404), for NK92 cells or primary NK cells respectively. Lipid preparations were infused into plasma-cleaned glass coverslips affixed within six-lane adhesive chambers (Ibidi, 80606) for 20 mins. SLBs were washed three times with 200 µL HBSS and blocked with 2 % BSA/HBSS for 20 min. The SLBs were washed again before protein dilutions were added for 20 mins to achieve the protein densities indicated in Table S3. Following this, the SLBs were washed. For those used to study primary NK cells, 2 µgmL^−1^ anti-RH5 clone R5.016 Alexa Fluor 568 was added for 20 mins before washing.

Primary NK cells or NK92 cells were prepared at a density of 5×10^6^ cells mL^−1^ in HBSS and warmed to 37 °C. In studies in which antibodies were used, 10 µg mL^−1^ W6/32 mIgG2a (Biolegend, 311441) was added prior to warming the cells. 100 µL of cell suspension was added to each SLB lane and was incubated at 37 °C for 30 mins. The SLBs were then fixed in 4 % paraformaldehyde (PFA, ThermoFisher, 28908) in HBSS for 10 mins, followed by washing. The cells were permeabilised using 0.1 % Triton-X100 (Sigma, T9284) in HBSS for 4 minutes before blocking with 5 % BSA/HBSS. Perforin staining was performed using 10 µg mL^−1^ anti-perforin (clone: δG9) Alexa Fluor 488 (Biolegend, 308108), for primary NK cells, or 10 µg mL^−1^ anti-perforin (clone: δG9) Alexa Fluor 594 (Biolegend, 308124) for NK92 cells. The antibody was incubated for either 1 hour at RT or overnight at 4 °C. SLB were washed three times prior to imaging. Imaging was performed using an Olympus cell TIRF 4Line system with a x100 (NA:1.45) oil objective (Olympus) at RT. Image analysis was performed using imageJ (FIJI) v1.54p. For TIRF imaging, cells were segmented on IRM prior to calculating average fluorescent intensity.

### Structured Illumination Microscopy (SIM)

SLBs were prepared for NK92 cells as previously stated. After fixation, cells were permeabilised with 4 % PFA v/v in HBSS for 4 mins prior to blocking with 5 % BSA w/v in HBSS for 1 hour and then 10 % v/v mouse serum in HBSS for 1 hour. The cells were then stained with a 1:400 dilution of ChromoTek GFP-Booster Alexa Fluor 647 (Proteintech, gb2AF647) for 20 mins at RT and were washed three times before imaging. Imaging was performed on a Zeiss Elyra 7 Lattice SIM microscope running Zeiss Zen Black v3.0, using a Plan-Apochromat 63x 1.4 NA Oil objective. SIM imaging using a 642 nm laser was performed on the basal membrane of the NK92 cells. SIM, using standard settings, and WF images were reconstructed using the Zeiss Zen Black v3.0 software. Following reconstruction, images were analysed in imageJ (FIJI) v1.54p. Cells were segmented on the generated WF image, before a 0.8 scale was used to exclude the edge of the membrane. The resulting ROI was used to identify LILRB1 clusters above a fluorescence intensity of 3000. Median integrated density was calculated for each cell.

### FRET microscopy

1×10^6^ HEK WT cells or HEK B2M^−/-^ cells were plated into one well of a 6 well plate on day 0. 2 µg of FRET reporter and Lck-mCherry2 were transfected into HEK cells on day 1. For each transfection, 4 µL of GeneJuice transfection reagent was added to 100 µL of antibiotic free DMEM, and then vortexed for 30 seconds. This was incubated at RT for 5 mins. 2 µg of PB-LILRB1^WT^ FRET or PB-LILRB1^BUCLD^ FRET and 2 µg of pHR-Lck-mCherry2 were added to each reaction and this was mixed by brief pipetting. The transfection mixture was incubated at room temperature for 15 mins before being added to the HEK cells in a dropwise manner. HEK cells were then harvested on day 4 and washed once in PBS before resuspending in cDMEM. Ibidi 8-well µ-slides were coated with 20% v/v Poly-L-Lysine (PLL, P4707) in PBS for 20 mins at room temperature. The PLL was aspirated and wells were washed with excess PBS three times. 6×10^5^ HEK cells were added to each well for 3 hours to adhere. In conditions where antibodies were used, 10 µg mL^−1^ W6/32 mIgG2a was added to the well with the cells. After 3 hours, media was aspirated and replaced with warmed 4% PFA v/v in PBS for 20 mins at 37 °C. The PFA was then aspirated and the wells were washed with 200 µL PBS three times and then left in 200 µL PBS for imaging.

Imaging was performed on a Zeiss Confocal 980 microscope, running Zeiss Zen Blue v3.3 using a Plan-Apochromat 63x/1.40 NA oil objective in confocal unidirectional line scanning mode. A 440 nm laser was used to excite both the CFP (detection wavelength: 464-499 nm) and Y-Pet (detection wavelength: 517-579 nm) for a series of 5 images of the basal membrane of the cell. Fluorophore emission detected by a 32 channel GaAsP-PMT detector, with CFP detection wavelength set to 464-499 nm and Y-PET detection wavelength set to 517-579 nm. A region of interest was then photobleached by scanning with a 514 nm laser 25 times. Following photobleaching, another series of 5 images was taken using the 440 nm laser. Images were analysed using imageJ (FIJI) v1.54p. Cells were first segmented on CFP fluorescent intensity; with regions of interest imported with image metadata. Mean fluorescence intensities for the CFP and Y-PET channels were then calculated for the whole cell and the region of interest, before comparing fluorescence intensity of the region of interest to the rest on the cell.

### Scanning Fluorescence Correlation Spectroscopy (FCS)

This was performed as previously described ^59,60^. In brief, HEK β2M^−/-^ cells were plated at 1×10^5^ cells per well of a PLL-coated 8-well Ibidi µ-slide on day 0. On day 1, these cells were transfected with pCDNA3.1(+)-HLA-A2-9V SCT mScarlet3 and either pHR-LILRB1^WT^-GFP or pHR-LILRB1^BUCLD^-GFP; as described previously. The following day, where indicated, 10 µg mL^−1^ of either the W6/32 mAb or RIFIN was added at 37 °C, 30 mins prior to imaging. HEK β2M-/- cells were imaged on a Zeiss LSM980 using a C-Apochromat 40 x 1.20 NA water immersion objective with 200,000 frames of a 128 px line at the basal cell surface unidirectionally scanned with the 488nm and 561 nm lasers using a pixel time of 0.98 µs and a pinhole size of 37 µm (1 AU equivalent). Raw data processing was performed using FoCuS-scan v1_15_107, followed by curve fitting using PyCorrFit v1.1.7.

### Statistical analysis

GraphPad Prism v10 was used for statistical analysis. Ordinary one-way analysis of variance (ANOVA) with Tukey’s post-hoc multiple comparison test. Where error bars are shown, they indicate SEM or SD as detailed in the figure legends. *p < 0.05, **p < 0.01, ***p < 0.001, ****p < 0.0001.

## Acknowledgements

This work was funded by a Wellcome Trust collaborative Award (224343/Z/21/Z), Japan Agency for Medical Research and Development (JP223fa627002), Japan Science and Technology (CREST, JPMJCR23B5), Japan Society for the Promotion of Science (24K2205, 23K27401) and Kennedy Trust for Rheumatology Research Cell Dynamics Platform. This work was supported by the Chinese Academy of Medical Sciences (CAMS) Innovation Fund for Medical Science (CIFMS), China (grant number: 2024-I2M-2-001-1). In addition, this work was supported by the Nippon Foundation and Shionogi Infectious Disease Research Promotion Foundation. We gratefully acknowledge the Oxford-ZEISS Centre of Excellence (Oxford-ZEISS CoE) in Biomedical Imaging, especially Dr. Kseniya Korobchevskaya and Dr. Christoffer Lagerholm, for their support and assistance in this work. The Oxford-ZEISS CoE is supported by the Kennedy Trust for Rheumatology Research, IDRM and Carl Zeiss GMBH. We thank Hannah Ivison for lab management. The authors would like to thank Dr Ed Lowe and the beamlines scientists at Diamond beamline I04 for help with crystallographic data collection.

## Author Contributions

S.G.C., M.A.W., S.I., M.D. and M.K.H. designed and conceived the study and wrote the manuscript. S.G.C. produced and tested RIFINs from African strains and conducted all other structural and biophysical analysis. S.I. and A.S. created RIFIN-expression libraries and identified LILRB1-D3D4-binding RIFINs. M.A.W., and A.M.M. performed SLB assay and immune cell studies. R.S, E.K., L.C. and S.V. contributed reagents.

## Data availability

Data within graphs (source data) are included with this submission. Crystallographic data is deposited in the protein data bank with accession codes 29WH and 29WI. Sequence data related to rif-lib1 and -lib2 were deposited at NCBI Gene expression omnibus with accession number GSE326451. All materials are available from the authors.

## Supplementary Figures

**Supplementary Figure 1:**
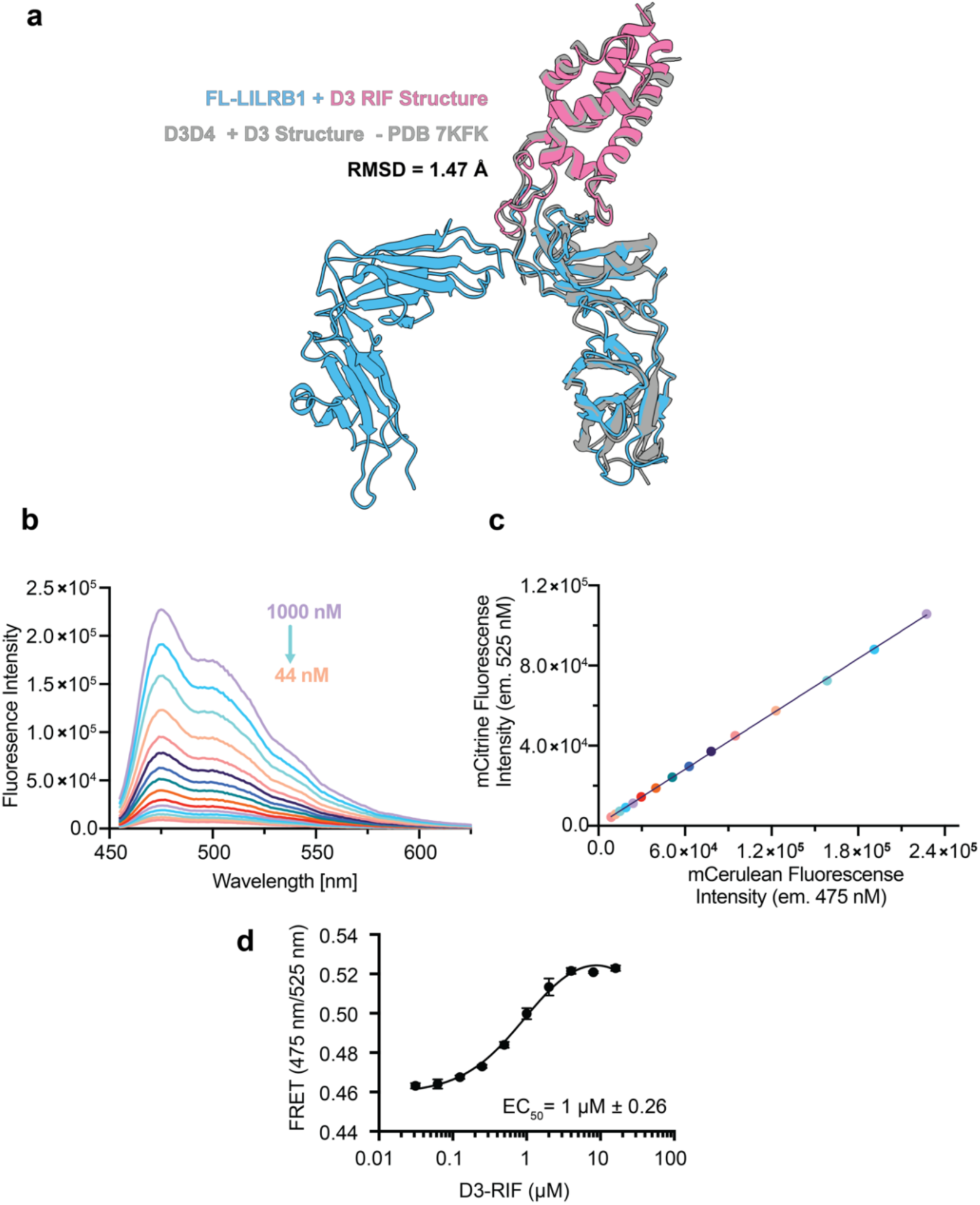
**a)** Structural overlay between the previously solved structure of D3-RIF bound to LILRB1-D3D4 (PDB accession code 7KFK, grey) and the crystal structure presented in this paper with the same RIFIN bound to full-length LILRB1 (pink/blue). The RMSD across all shared backbone atoms was measured to be 1.47 Å. **b)** Emission spectrum of the LILRB1 FRET-probe between 460 and 625 nm after excitation at 433 nm recorded at probe concentrations from 44 nM to 1000 nM, n=1. **c)** Plotting the fluorescence intensity of mCitrine and mCerulean across the concentration ranges described above. Any deviation from a straight line would indicate a concentration dependent FRET effect, n =1. **d)** FRET ratio of 1 µM of LILRB1 FRET-probe when titrated against an increasing concentration of D3-RIF. EC50 value was calculated in PRISM using the one-site total model, n=3.

**Supplementary Figure 2:**
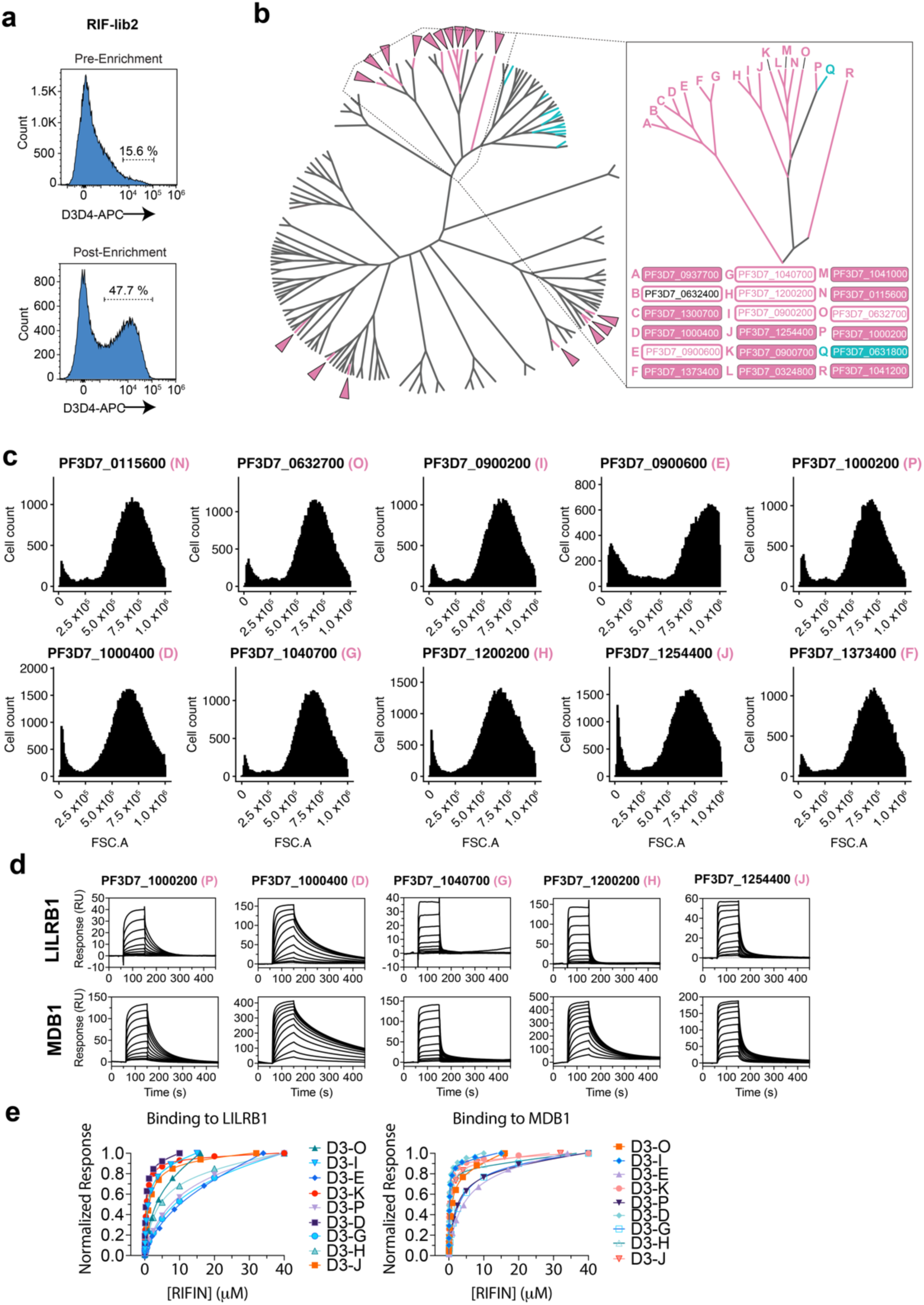
**a)** Pre and post enrichment flow sorting with LILRB1-D3D4-Fc for rif-lib2 (biological duplicate). **b)** Full 3D7 RIFIN phylogenetic tree (left) and annotated region of the major LILRB1-D3D4 binding clade (right) as shown in Figure 2b and c. **b)** Assessment of iRBC transfected with candidate RIFINs binding to fluorescent LILRB1-D3D4-Fc. **c)** Sensorgrams showing candidate RIFINs binding to both full-length LILRB1 (top) and D3D4-containing MDB1 antibody (bottom), data from three independent experiments. **d)** Steady state affinity fits of LILRB1 (left) and MDB1 (right) with candidate RIFINs. Data is fitted using the one-site-total PRISM model, n =3.

**Supplementary Figure 3:**
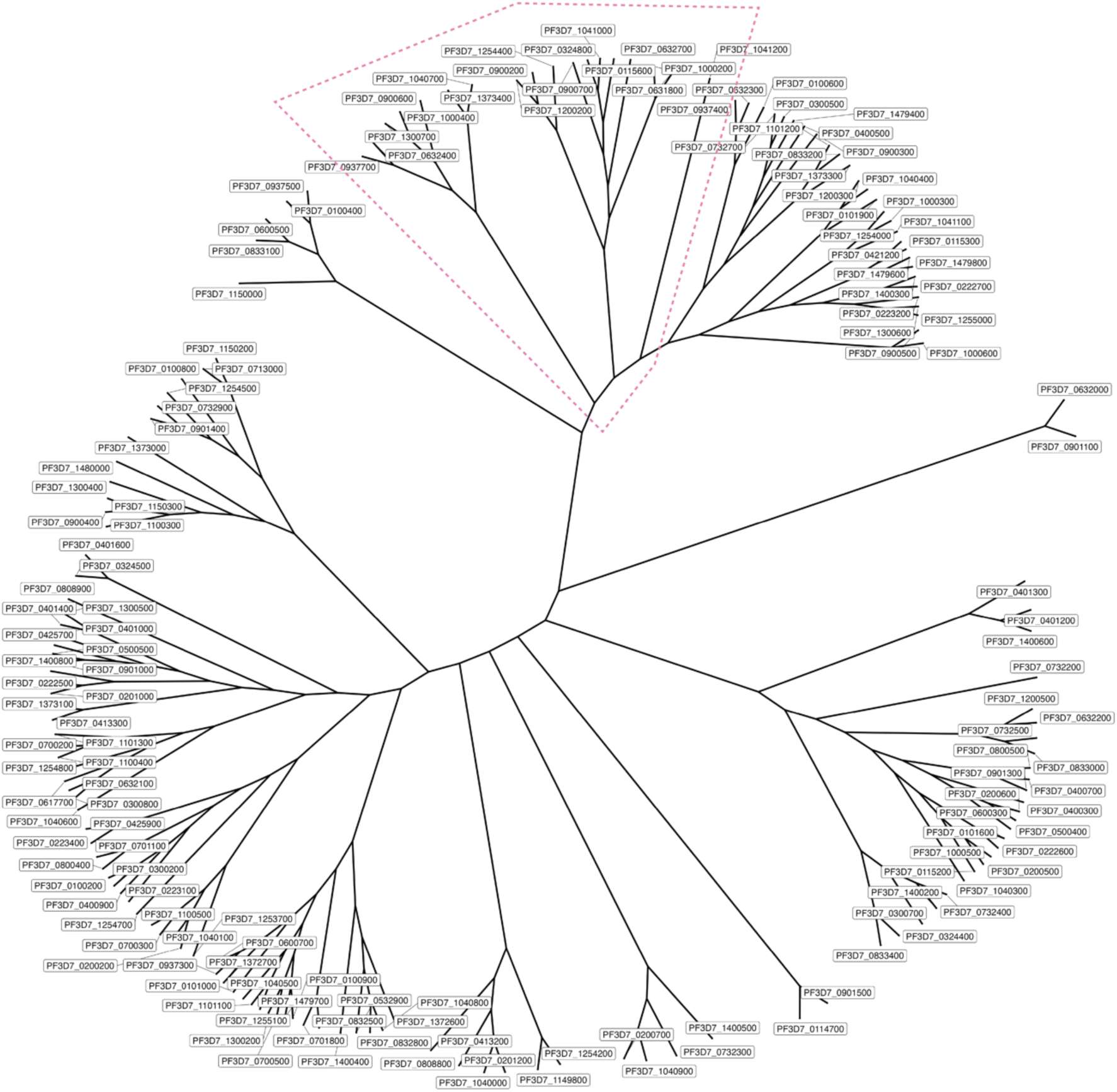
Fully annotated phylogenetic tree of every RIFIN of the *Pf*3D7 genome. The region shown in Figure 2 c is highlighted within the pink dashed line. The tree was calculated using Clustal Omega and plotted using the ggtree R package^57^.

**Supplementary Figure 4:**
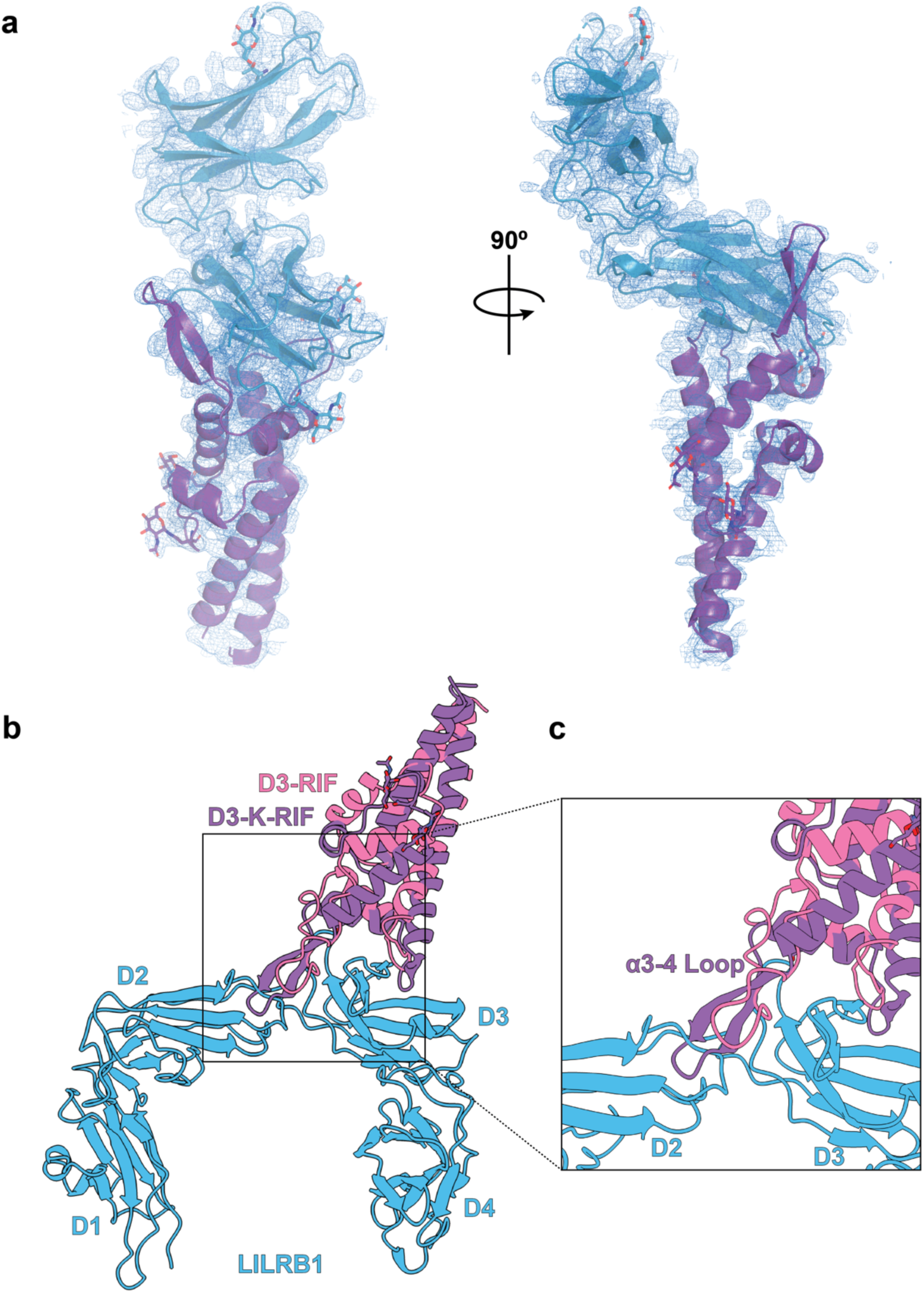
**a)** Cartoon representation of D3-K-RIF (purple) bound to LILRB1-D3D4 (blue) showing the electron density as a mesh. PDB accession 29WH. **b)** Overlay showing a superimposition of D3-RIF (pink) and D3-K (purple) bound to LILRB1 (blue). **c)** Zoomed region showing that with this superimposition, a clash between the ⍺3-4 binding loop of the RIFIN and domain 2 of LILRB1 is observed.

**Supplementary Figure 5:**
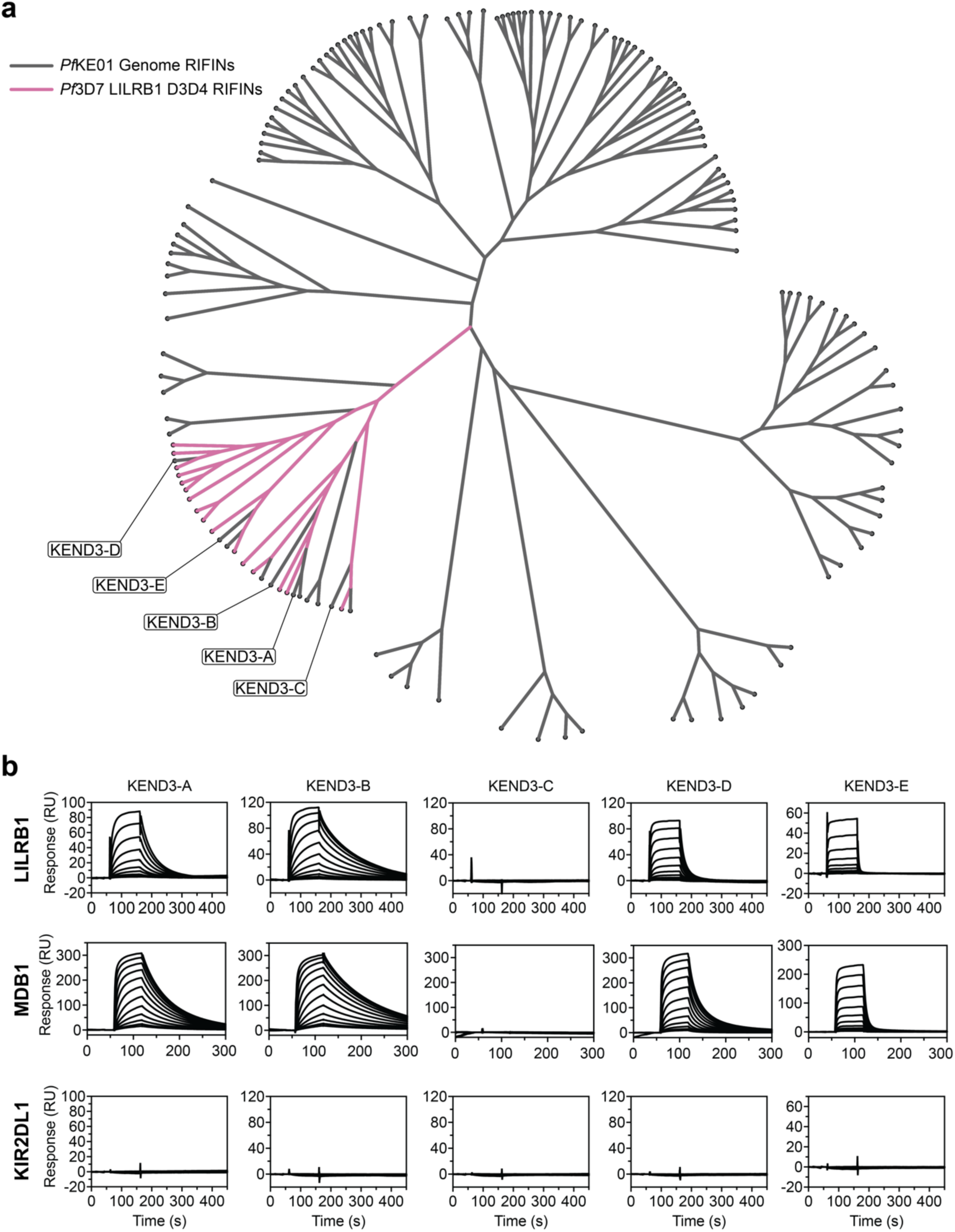
**a)** Phylogenetic tree of all RIFINs from the Kenyan strain *Pf*KE01 (grey) with the confirmed LILRB1-D3D4-binding RIFINs of *Pf*3D7 (pink). Annotated are 5 RIFINs that were predicted to bind LILRB1-D3D4 based on their phylogenetic similarity to 3D7 LILRB1-D3D4-binding RIFINs and that expressed in sufficient quantities for SPR analysis. **b)** SPR sensorgrams showing the response when RIFINs are flowed over either full-length LILRB1 (top row), MDB1 (middle row) or KIR2DL1 as a negative control (bottom row), n=3.

**Supplementary Figure 6.**
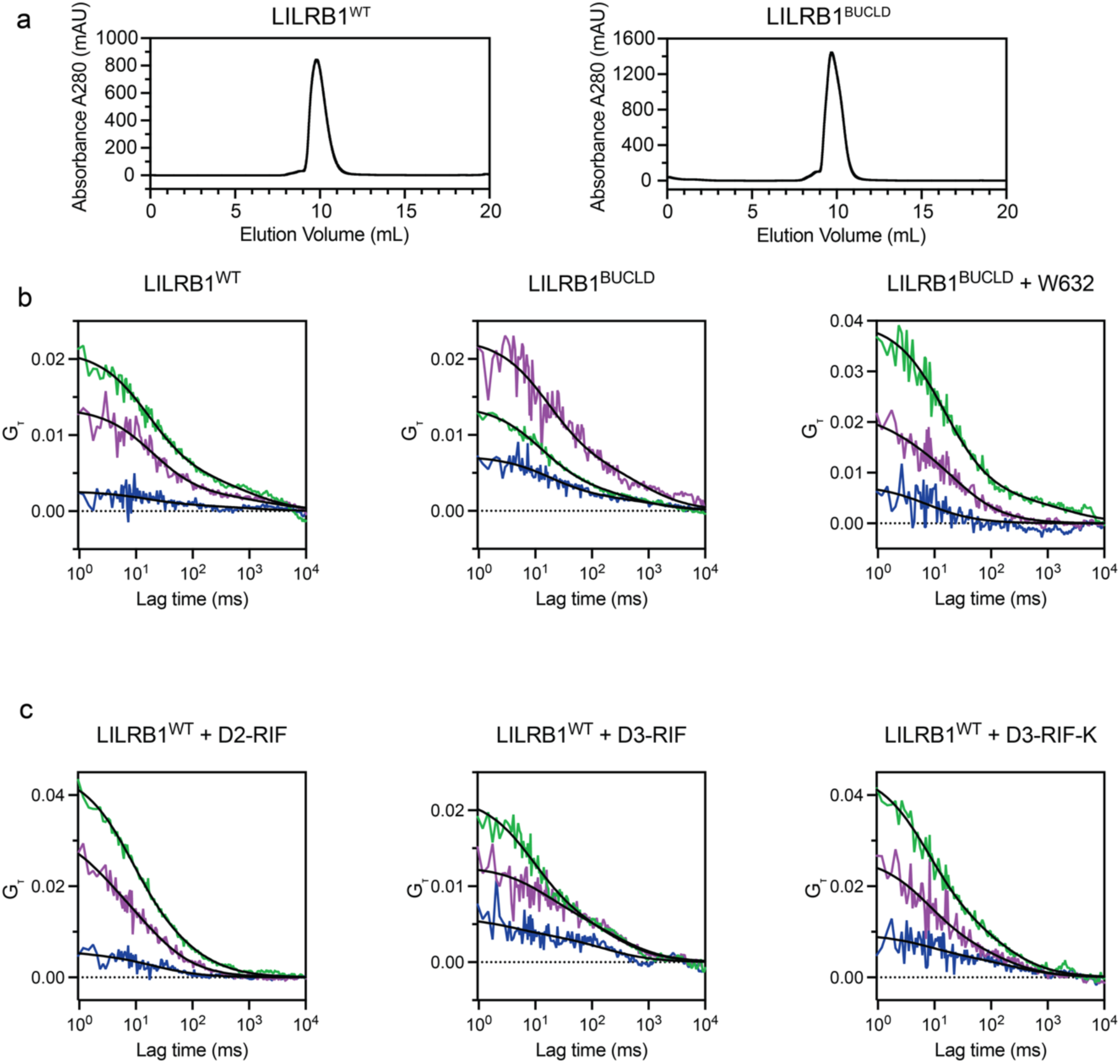
**a):** Size exclusion chromatography chromatograms showing the elution profile of LILRB1^WT^ (left) and LILRB1^BUCLD^ (right). **a)** and **b)** Representative FCS and FCCS curves from the specified conditions. Green line represents LILRB1-GFP, magenta line represents HLA-A2-9V SCT mScarlet3, blue line represents cross-correlation, and black lines show fitting curves. Representative figures from three independent experiments consisting of: LILRB1^WT^: 21 cells; LILRB1^BUCLD^: 27 cells; LILRB1^BUCLD^ + W6/32: 27 cells; LILRB1^WT^ + D2-RIF: 20 cells; LILRB1^WT^ + D3-RIF: 27 cells; LILRB1^WT^ + D3-K-RIF: 23 cells.

**Supplementary Figure 7:**
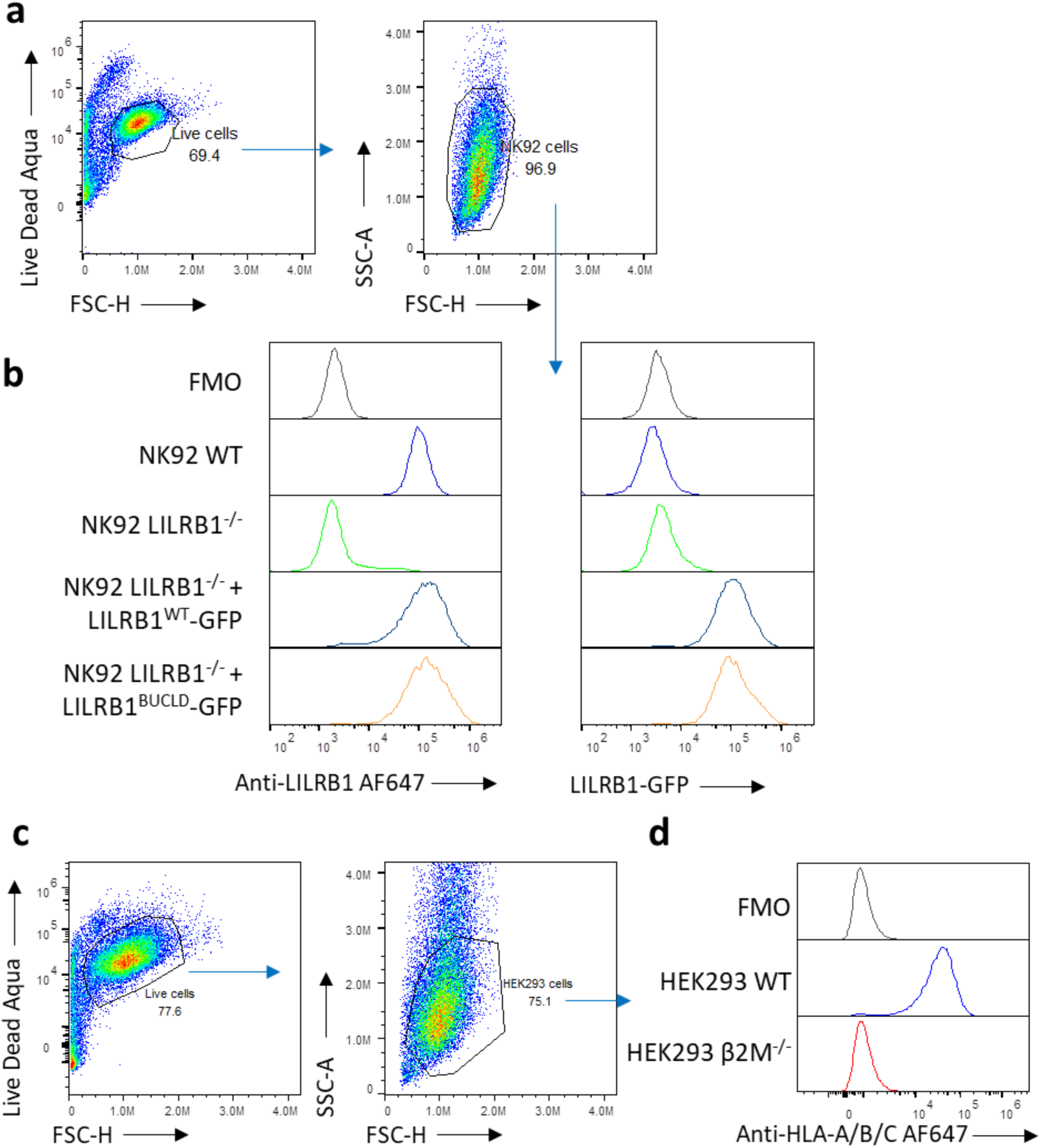
**a)** Gating strategy to assess LILRB1 expression on NK92 cells. **b)** Expression of LILRB1 on NK92 cells through either anti-LILRB1 staining (left) or GFP expression (right). **c)** Gating strategy to assess expression of HLA-A/B/C on HEK293 cells. **d)** Expression of HLA-A/B/C on HEK293 WT or HEK293 β2M^−/-^ cells.

**Supplementary Figure 8:**
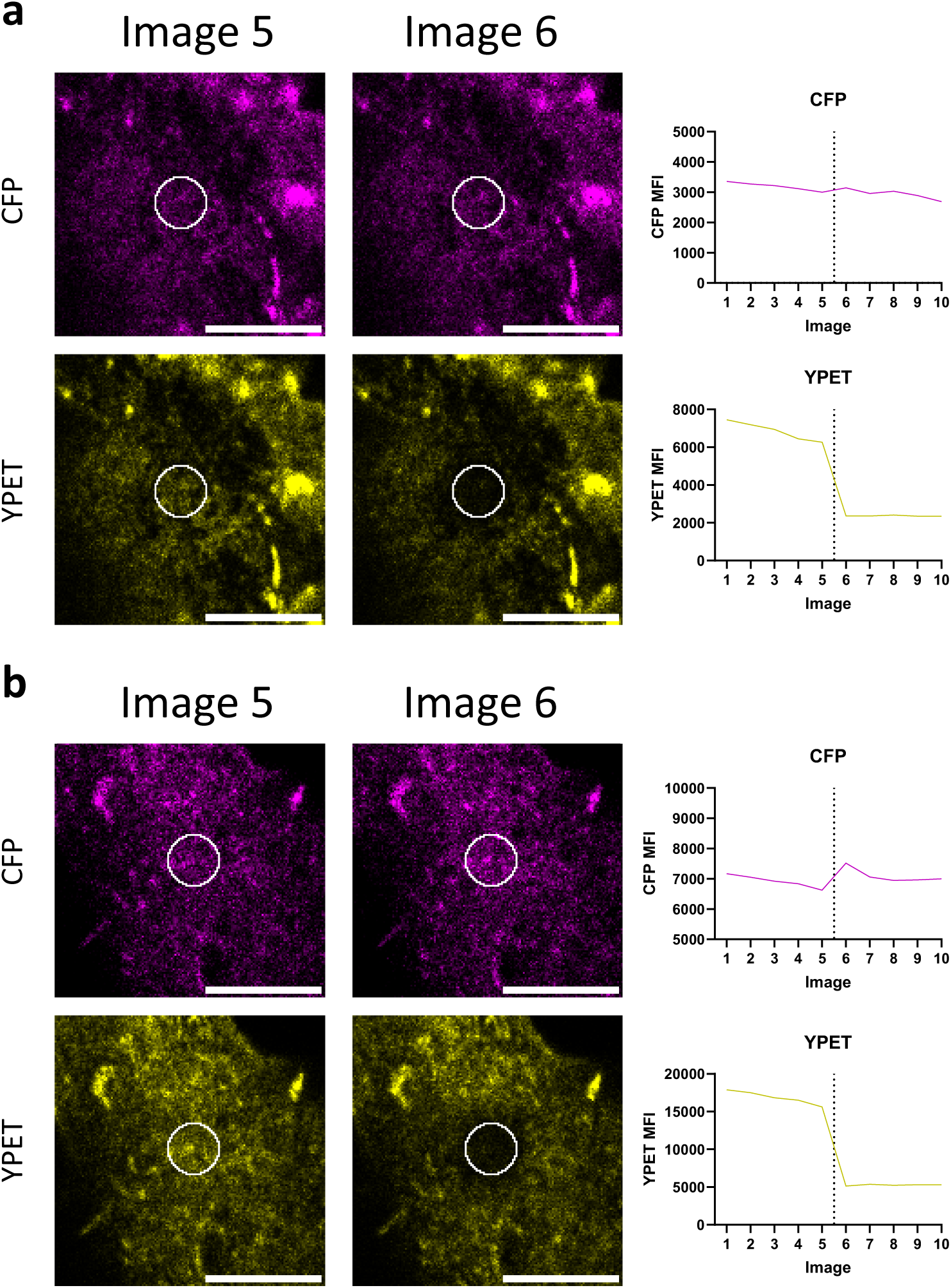
Representative images of **a)** LILRB1^WT^ or **b)** LILRB1^BUCLD^ FRET reporter, showing image 5 prior to photobleaching (left); image 6 post photobleaching (centre) and mean fluorescence from the region of interest indicated in white in each image (right). Representative figures from three independent experiments consisting of: LIILRB1^WT^: 45 cells; LILRB1^BUCLD^: 45 cells; LILRB1^BUCLD^ + W6/32: 39 cells.

**Supplementary Figure 9:**
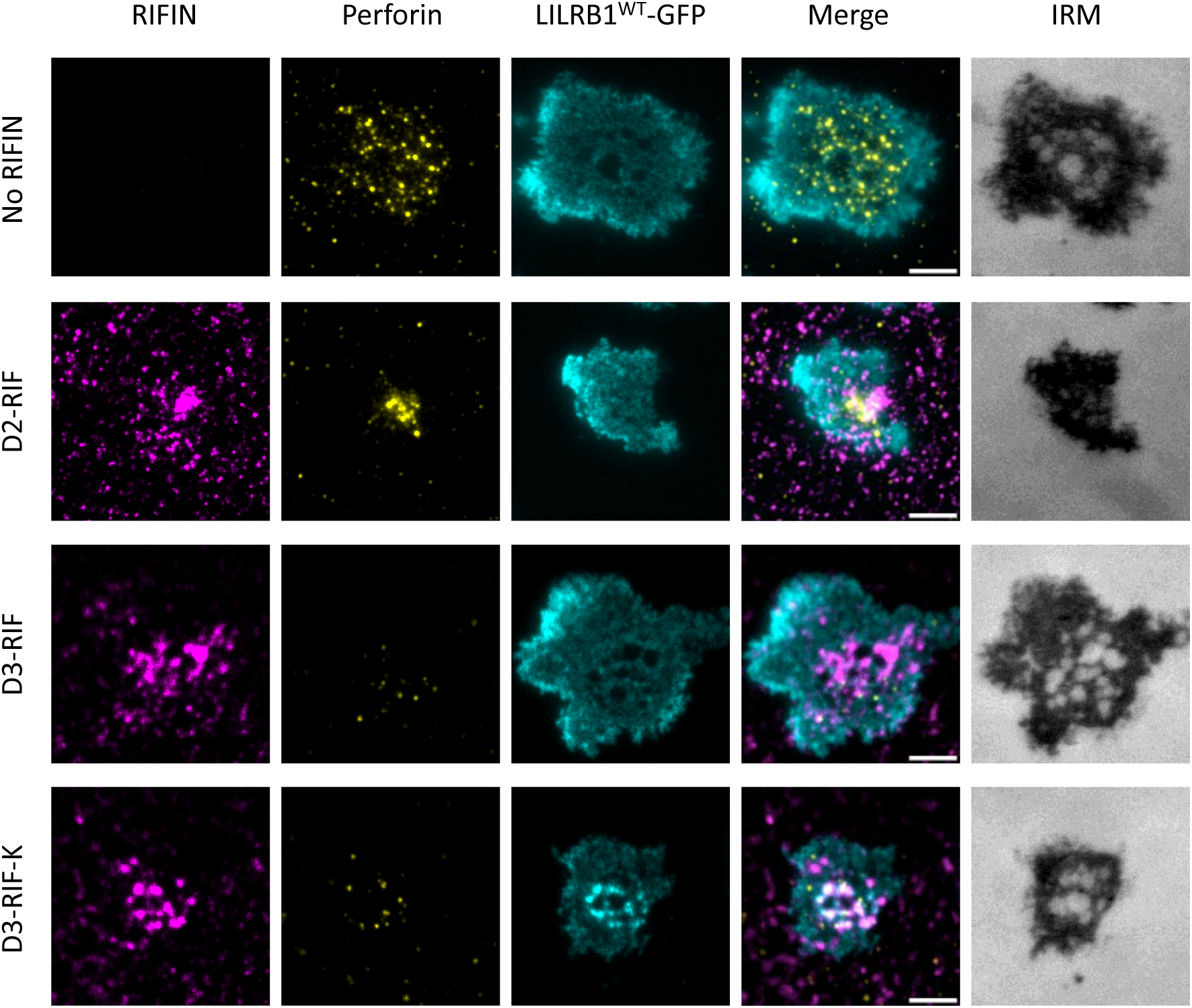
Representative images from the contact area of NK92 cells expressing LILRB1^WT^-GFP when added to activating supported lipid bilayers coated with either no RIFIN, D2-RIF, D3-RIF or D3-K-RIF for 30 mins. Scale bars 5 µm.

**Supplementary Table 1.**
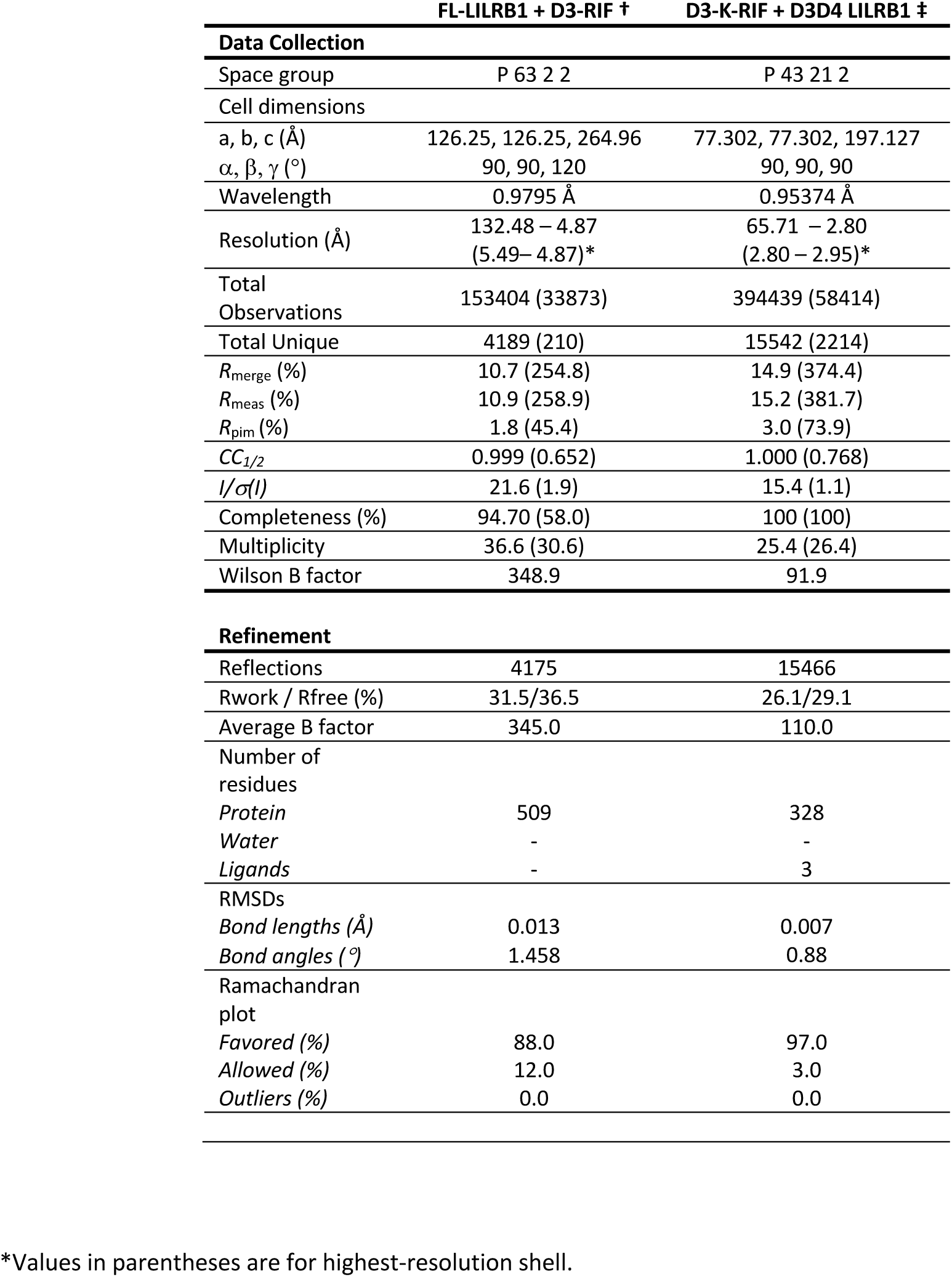
Crystallographic data collection and refinement statistics.

**Supplementary Table 2:** Construct boundaries and mutations for expressed RIFINs.

| Accession Number | RIFIN Name | Domain Boundaries | Mutations |
| --- | --- | --- | --- |
| *Pf*3D7_1254800 | D2-RIF | 165 – 274 | E179C, C223S, A263C |
| *Pf*3D7_1373400 | D3-RIF | 173 – 326 | N/A |
| *Pf*3D7_1000400 | D3-D-RIF | 179 – 320 | G191C, A296C |
| *Pf*3D7_0900600 | D3-E-RIF | 173 – 327 | G183C, A296C |
| *Pf*3D7_1040700 | D3-G-RIF | 176 – 322 | A193C, A297C |
| *Pf*3D7_1200200 | D3-H-RIF | 182 – 327 | A182C, L137C |
| *Pf*3D7_0900200 | D3-I-RIF | 180 – 325 | A182C, V316C |
| *Pf*3D7_1254400 | D3-J-RIF | 178 – 328 | G184C, E315C |
| *Pf*3D7_0900700 | D3-K-RIF | 178 – 331 | G185C, T317C |
| *Pf*3D7_0632700 | D3-O-RIF | 180 – 324 | G187C, A308C |
| *Pf*3D7_1000200 | D3-P-RIF | 172 – 320 | G179C, A307C |
| *Pf*KE01_080005900 | KEND3-A | 185-309 | A190C, V304C |
| *Pf*KE01_040007100 | KEND3-B | 177-314 | G183C, K307C |
| *Pf*KE01_000005700 | KEND3-C | 152-292 | G160C, A283C |
| *Pf*KE01_120005500 | KEND3-D | 175-321 | G180C, A312C |
| *Pf*KE01_040032500 | KEND3-E | 179-305 | G189C, A293C |

**Supplementary Table 3:** Protein densities for supported lipid bilayers.

| NK92 cells | Molecules $\mu\text{m}^{-2}$ |
| --- | --- |
| ICAM1 | 1000 |
| B7-H6 | 200 |
| CD58 | 200 |
| D2 Rif | 100 |
| D3 Rif | 100 |
| D3-K Rif | 100 |
| Primary NK cells | Molecules $\mu\text{m}^{-2}$ |
| ICAM1 | 200 |
| RH5 | 100 |
| D2 Rif | 100 |
| D3 Rif | 100 |
| D3-K Rif | 100 |

## References

1. Moreau, A., Yaya, F., Lu, H., Surendranath, A., Charrier, A., Dehapiot, B., Helfer, E., Viallat, A., and Peng, Z. (2023). Physical mechanisms of red blood cell splenic filtration. Proc. Natl. Acad. Sci. U. S. A. 120. 10.1073/PNAS.2300095120/FORMAT/EPUB.

2. Chamberlain, S.G., Iwanaga, S., and Higgins, M.K. (2025). Immune evasion runs in the family: two surface protein families of Plasmodium falciparum–infected erythrocytes. Curr. Opin. Microbiol. 85, 102598. 10.1016/J.MIB.2025.102598.

3. Chan, J.A., Boyle, M.J., Moore, K.A., Reiling, L., Lin, Z., Hasang, W., Avril, M., Manning, L., Mueller, I., Laman, M., et al. (2019). Antibody Targets on the Surface of Plasmodium falciparum–Infected Erythrocytes That Are Associated With Immunity to Severe Malaria in Young Children. J. Infect. Dis. 219, 819–828. 10.1093/INFDIS/JIY580.

4. Bull, P.C., Lowe, B.S., Kortok, M., Molyneux, C.S., Newbold, C.I., and Marsh, K. (1998). Parasite antigens on the infected red cell surface are targets for naturally acquired immunity to malaria. Nature Medicine 1998 4:3 *4*, 358–360. 10.1038/nm0398-358.

5. Hart, G.T., Tran, T.M., Theorell, J., Schlums, H., Arora, G., Rajagopalan, S., Jules Sangala, A.D., Welsh, K.J., Traore, B., Pierce, S.K., et al. (2019). Adaptive NK cells in people exposed to Plasmodium falciparum correlate with protection from malaria. Journal of Experimental Medicine 216. 10.1084/jem.20181681.

6. Arora, G., Hart, G.T., Manzella-Lapeira, J., Doritchamou, J.Y.A., Narum, D.L., Thomas, L.M., Brzostowski, J., Rajagopalan, S., Doumbo, O.K., Traore, B., et al. (2018). NK cells inhibit plasmodium falciparum growth in red blood cells via antibody-dependent cellular cytotoxicity. Elife 7. 10.7554/eLife.36806.

7. Junqueira, C., Polidoro, R.B., Castro, G., Absalon, S., Liang, Z., Sen Santara, S., Crespo, Â., Pereira, D.B., Gazzinelli, R.T., Dvorin, J.D., et al. (2021). γδ T cells suppress Plasmodium falciparum blood-stage infection by direct killing and phagocytosis. Nat. Immunol. 22, 347–357. 10.1038/s41590-020-00847-4.

8. Saito, F., Hirayasu, K., Satoh, T., Wang, C.W., Lusingu, J., Arimori, T., Shida, K., Palacpac, N.M.Q., Itagaki, S., Iwanaga, S., et al. (2017). Immune evasion of Plasmodium falciparum by RIFIN via inhibitory receptors. Nature 2017 552:7683 *552*, 101–105. 10.1038/nature24994.

9. Sakoguchi, A., Saito, F., Hirayasu, K., Shida, K., Matsuoka, S., Itagaki, S., Nakai, W., Kohyama, M., Suenaga, T., Iwanaga, S., et al. (2021). Plasmodium falciparum RIFIN is a novel ligand for inhibitory immune receptor LILRB2. Biochem. Biophys. Res. Commun. 548, 167–173. 10.1016/J.BBRC.2021.02.033.

10. Xie, Y., Li, X., Chai, Y., Song, H., Qi, J., and Gao, G.F. (2021). Structural basis of malarial parasite RIFIN-mediated immune escape against LAIR1. Cell Rep. 36. 10.1016/j.celrep.2021.109600.

11. Sakoguchi, A., Chamberlain, S.G., Mørch, A.M., Widdess, M., Harrison, T.E., Dustin, M.L., Arase, H., Higgins, M.K., and Iwanaga, S. (2025). RIFINs displayed on malaria-infected erythrocytes bind KIR2DL1 and KIR2DS1. Nature 2025 643:8074 *643*, 1363– 1371. 10.1038/s41586-025-09091-y.

12. Chen, Y., Xu, K., Piccoli, L., Foglierini, M., Tan, J., Jin, W., Gorman, J., Tsybovsky, Y., Zhang, B., Traore, B., et al. (2021). Structural basis of malaria RIFIN binding by LILRB1-containing antibodies. Nature 2021 592:7855 *592*, 639–643. 10.1038/s41586-021-03378-6.

13. Chamberlain, S.G., Iwanaga, S., and Higgins, M.K. (2025). Immune evasion runs in the family: two surface protein families of Plasmodium falciparum–infected erythrocytes. Curr. Opin. Microbiol. 85, 102598. 10.1016/J.MIB.2025.102598.

14. Cuapio, A., and Ljunggren, H.G. (2025). Both inhibitory and activating KIRs recognize RIFINs: a dual-edged mechanism of NK cell control in malaria. Signal Transduction and Targeted Therapy 2025 10:1 *10*, 253-. 10.1038/s41392-025-02341-5.

15. Cosman, D., Fanger, N., Borges, L., Kubin, M., Chin, W., Peterson, L., and Hsu, M.L. (1997). A novel immunoglobulin superfamily receptor for cellular and viral MHC class I molecules. Immunity 7, 273–282. 10.1016/S1074-7613(00)80529-4.

16. Colonna, M., Navarro, F., Bellón, T., Llano, M., García, P., Samaridis, J., Angman, L., Cella, M., and López-Botet, M. (1997). A Common Inhibitory Receptor for Major Histocompatibility Complex Class I Molecules on Human Lymphoid and Myelomonocytic Cells. Journal of Experimental Medicine 186, 1809–1818. 10.1084/JEM.186.11.1809.

17. Harrison, T.E., Mørch, A.M., Felce, J.H., Sakoguchi, A., Reid, A.J., Arase, H., Dustin, M.L., and Higgins, M.K. (2020). Structural basis for RIFIN-mediated activation of LILRB1 in malaria. Nature 2020 587:7833 *587*, 309–312. 10.1038/s41586-020-2530-3.

18. Wang, Q., Song, H., Cheng, H., Qi, J., Nam, G., Tan, S., Wang, J., Fang, M., Shi, Y., Tian, Z., et al. (2020). Structures of the four Ig-like domain LILRB2 and the four-domain LILRB1 and HLA-G1 complex. Cell. Mol. Immunol. 17, 966–975. 10.1038/s41423-019-0258-5.

19. Kovalevskiy, O., Nicholls, R.A., and Murshudov, G.N. (2016). Automated refinement of macromolecular structures at low resolution using prior information. urn:issn:2059-7983 72, 1149–1161. 10.1107/S2059798316014534.

20. Mohammed, F., Stones, D.H., and Willcox, B.E. (2019). Application of the immunoregulatory receptor LILRB1 as a crystallisation chaperone for human class I MHC complexes. J. Immunol. Methods 464, 47–56. 10.1016/J.JIM.2018.10.011.

21. Bacia, K., Kim, S.A., and Schwille, P. (2006). Fluorescence cross-correlation spectroscopy in living cells. Nature Methods 2006 3:2 *3*, 83–89. 10.1038/nmeth822.

22. Yu, Y.Y.L., Netuschil, N., Lybarger, L., Connolly, J.M., and Hansen, T.H. (2002). Cutting Edge: Single-Chain Trimers of MHC Class I Molecules Form Stable Structures That Potently Stimulate Antigen-Specific T Cells and B Cells. The Journal of Immunology 168, 3145–3149. 10.4049/JIMMUNOL.168.7.3145.

23. Mørch, A.M., Schneider, F., Jenkins, E., Santos, A.M., Fraser, S.E., Davis, S.J., and Dustin, M.L. (2022). The kinase occupancy of T cell coreceptors reconsidered. Proc. Natl. Acad. Sci. U. S. A. 119, e2213538119–e2213538119. 10.1073/pnas.2213538119.

24. Cyster, J.G., and Goodnow, C.C. (1997). Tuning antigen receptor signaling by CD22: Integrating cues from antigens and the microenvironment. Immunity 6, 509–517. 10.1016/S1074-7613(00)80339-8.

25. Zhao, Y., Harrison, D.L., Song, Y., Ji, J., Huang, J., and Hui, E. (2018). Antigen-Presenting Cell-Intrinsic PD-1 Neutralizes PD-L1 in cis to Attenuate PD-1 Signaling in T Cells. Cell Rep. 24, 379–390.e6. 10.1016/j.celrep.2018.06.054.

26. Back, J., Chalifour, A., Scarpellino, L., and Held, W. (2007). Stable masking by H-2Dd cis ligand limits Ly49A relocalization to the site of NK cell/target cell contact. Proc. Natl. Acad. Sci. U. S. A. 104, 3978–3983. 10.1073/pnas.0607418104.

27. Held, W., and Mariuzza, R.A. (2011). Cis–trans interactions of cell surface receptors: biological roles and structural basis. Cell. Mol. Life Sci. 68, 3469. 10.1007/S00018-011-0798-Z.

28. Masuda, A., Nakamura, A., Maeda, T., Sakamoto, Y., and Takai, T. (2007). Cis binding between inhibitory receptors and MHC class I can regulate mast cell activation. Journal of Experimental Medicine 204, 907–920. 10.1084/JEM.20060631.

29. Hui, E. (2023). Cis Interactions of Membrane Receptors and Ligands. Annu. Rev. Cell Dev. Biol. 39, 391–408. 10.1146/ANNUREV-CELLBIO-120420-103941/CITE/REFWORKS.

30. Wang, H., Zheng, X., Wei, H., Tian, Z., and Sun, R. (2012). Important Role for NKp30 in Synapse Formation and Activation of NK Cells. Immunol. Invest. 41, 367–381. 10.3109/08820139.2011.632799.

31. Sun, J., Lei, L., Tsai, C.M., Wang, Y., Shi, Y., Ouyang, M., Lu, S., Seong, J., Kim, T.J., Wang, P., et al. (2017). Engineered proteins with sensing and activating modules for automated reprogramming of cellular functions. Nature Communications 2017 8:1 8, 477-. 10.1038/s41467-017-00569-6.

32. Chua, C.L.L., Brown, G., Hamilton, J.A., Rogerson, S., and Boeuf, P. (2013). Monocytes and macrophages in malaria: Protection or pathology? Trends Parasitol. 29, 26–34. 10.1016/j.pt.2012.10.002.

33. Gupta, A., Kalkal, M., and Das, J. (2025). Emerging Role of Splenic Macrophage in Malaria Pathogenesis and Immunity. Immunity, Inflammation and Disease 13, e70258. 10.1002/iid3.70258.

34. Ozarslan, N., Robinson, J.F., and Gaw, S.L. (2019). Circulating Monocytes, Tissue Macrophages, and Malaria. J. Trop. Med. 2019, 3720838. 10.1155/2019/3720838.

35. Tan, J., Pieper, K., Piccoli, L., Abdi, A., Foglierini, M., Geiger, R., Maria Tully, C., Jarrossay, D., Maina Ndungu, F., Wambua, J., et al. (2015). A LAIR1 insertion generates broadly reactive antibodies against malaria variant antigens. Nature 2015 529:7584 529, 105–109. 10.1038/nature16450.

36. Xu, K., Wang, Y., Shen, C.H., Chen, Y., Zhang, B., Liu, K., Tsybovsky, Y., Wang, S., Farney, S.K., Gorman, J., et al. (2021). Structural basis of LAIR1 targeting by polymorphic Plasmodium RIFINs. Nature Communications 2021 12:1 *12*, 4226-. 10.1038/s41467-021-24291-6.

37. Alanine, D.G.W., Quinkert, D., Kumarasingha, R., Mehmood, S., Donnellan, F.R., Minkah, N.K., Dadonaite, B., Diouf, A., Galaway, F., Silk, S.E., et al. (2019). Human Antibodies that Slow Erythrocyte Invasion Potentiate Malaria-Neutralizing Antibodies. Cell 178, 216. 10.1016/j.cell.2019.05.025.

38. Waithe, D., Schneider, F., Chojnacki, J., Clausen, M.P., Shrestha, D., de la Serna, J.B., and Eggeling, C. (2018). Optimized processing and analysis of conventional confocal microscopy generated scanning FCS data. Methods 140–141, 62–73. 10.1016/J.YMETH.2017.09.010.

39. Robertson, M., Cochran, K.J., Cameron, C., Le, J., Tantravahi, R., and Ritz, J. (1996). Characterization of a cell line, NKL, derived from an aggressive human natural killer cell leukemia. Exp. Hematol.

40. Müller, P., Schwille, P., and Weidemann, T. (2014). PyCorrFit—generic data evaluation for fluorescence correlation spectroscopy. Bioinformatics 30, 2532–2533. 10.1093/BIOINFORMATICS/BTU328.

41. Agirre, J., Atanasova, M., Bagdonas, H., Ballard, C.B., Baslé, A., Beilsten-Edmands, J., Borges, R.J., Brown, D.G., Burgos-Mármol, J.J., Berrisford, J.M., et al. (2023). The CCP4 suite: integrative software for macromolecular crystallography. Acta Crystallogr. D Struct. Biol. 79, 449–461. 10.1107/S2059798323003595.

42. Winter, G. (2010). Xia2: An expert system for macromolecular crystallography data reduction. J. Appl. Crystallogr. 43, 186–190. 10.1107/S0021889809045701;PAGE:STRING:ARTICLE/CHAPTER.

43. Shen, W., Le, S., Li, Y., and Hu, F. (2016). SeqKit: A Cross-Platform and Ultrafast Toolkit for FASTA/Q File Manipulation. PLoS One 11. 10.1371/JOURNAL.PONE.0163962.

44. Evans, P. (2006). Scaling and assessment of data quality. Acta Crystallogr. D Biol. Crystallogr. 62, 72–82. 10.1107/S0907444905036693.

45. Bolger, A.M., Lohse, M., and Usadel, B. (2014). Trimmomatic: a flexible trimmer for Illumina sequence data. Bioinformatics 30, 2114–2120. 10.1093/BIOINFORMATICS/BTU170.

46. McCoy, A.J., Grosse-Kunstleve, R.W., Adams, P.D., Winn, M.D., Storoni, L.C., and Read, R.J. (2007). Phaser crystallographic software. urn:issn:0021–8898 *40*, 658–674. 10.1107/S0021889807021206.

47. Langmead, B., and Salzberg, S.L. (2012). Fast gapped-read alignment with Bowtie 2. Nat. Methods 9, 357–359. 10.1038/NMETH.1923.

48. Emsley, P., Lohkamp, B., Scott, W.G., and Cowtan, K. (2010). Features and development of Coot. urn:issn:0907–4449 *66*, 486–501. 10.1107/S0907444910007493.

49. Liao, Y., Smyth, G.K., and Shi, W. (2014). FeatureCounts: An efficient general purpose program for assigning sequence reads to genomic features. Bioinformatics 30, 923–930. 10.1093/BIOINFORMATICS/BTT656.

50. Abramson, J., Adler, J., Dunger, J., Evans, R., Green, T., Pritzel, A., Ronneberger, O., Willmore, L., Ballard, A.J., Bambrick, J., et al. (2024). Accurate structure prediction of biomolecular interactions with AlphaFold 3. Nature 2024 630:8016 *630*, 493–500. 10.1038/s41586-024-07487-w.

51. Pettersen, E.F., Goddard, T.D., Huang, C.C., Couch, G.S., Greenblatt, D.M., Meng, E.C., and Ferrin, T.E. (2004). UCSF Chimera--a visualization system for exploratory research and analysis. J. Comput. Chem. 25, 1605–1612. 10.1002/JCC.20084.

52. Chen, Y., Xu, K., Piccoli, L., Foglierini, M., Tan, J., Jin, W., Gorman, J., Tsybovsky, Y., Zhang, B., Traore, B., et al. (2021). Structural basis of malaria RIFIN binding by LILRB1-containing antibodies. Nature, 1–5. 10.1038/s41586-021-03378-6.

53. Liebschner, D., Afonine, P. V., Baker, M.L., Bunkoczi, G., Chen, V.B., Croll, T.I., Hintze, B., Hung, L.W., Jain, S., McCoy, A.J., et al. (2019). Macromolecular structure determination using X-rays, neutrons and electrons: recent developments in Phenix. Acta Crystallogr. D Struct. Biol. 75, 861–877. 10.1107/S2059798319011471.

54. Bricogne G., Blanc E., Brandl M., Flensburg C., Keller P., Paciorek W., Roversi P, Sharff A., Smart O.S., Vonrhein C., et al. (2017). BUSTER v2.10. Preprint.

55. Smart, O.S., Womack, T.O., Flensburg, C., Keller, P., Paciorek, W., Sharff, A., Vonrhein, C., and Bricogne, G. (2012). Exploiting structure similarity in refinement: Automated NCS and target-structure restraints in BUSTER. Acta Crystallogr. D Biol. Crystallogr. 68, 368–380. 10.1107/S0907444911056058/BA5178SUP1.PDF.

56. Boersma, A.J., Liu, B., Poolman, B., Eggers, D.K., Le, J.M., Pham, D.N., Nham, N.T., Contreras, F.A., Gnutt, D., Gao, M., et al. (2015). A sensor for quantification of macromolecular crowding in living cells. cell.com 108, 114a. 10.1016/j.bpj.2014.11.640.

57. Yu, G., Lam, T.T.Y., Zhu, H., and Guan, Y. (2018). Two Methods for Mapping and Visualizing Associated Data on Phylogeny Using Ggtree. Mol. Biol. Evol. 35, 3041–3043. 10.1093/molbev/msy194.

58. Sušac, L., Vuong, M.T., Thomas, C., von Bülow, S., O’Brien-Ball, C., Santos, A.M., Fernandes, R.A., Hummer, G., Tampé, R., and Davis, S.J. (2022). Structure of a fully assembled tumor-specific T cell receptor ligated by pMHC. Cell 185, 3201–3213.e19. 10.1016/j.cell.2022.07.010.

59. Philips, E.A., Liu, J., Kvalvaag, A., Mørch, A.M., Tocheva, A.S., Ng, C., Liang, H., Ahearn, I.M., Pan, R., Luo, C.C., et al. (2024). Transmembrane domain–driven PD-1 dimers mediate T cell inhibition. Sci. Immunol. 9. 10.1126/sciimmunol.ade6256.

60. Mørch, A.M., and Schneider, F. (2023). Investigating Diffusion Dynamics and Interactions with Scanning Fluorescence Correlation Spectroscopy (sFCS). Methods in Molecular Biology 2654, 61–89. 10.1007/978-1-0716-3135-5_5/TABLES/1.

